# TRIM67 Regulates Exocytic Mode and Neuronal Morphogenesis via SNAP47

**DOI:** 10.1101/2020.02.01.930404

**Authors:** Fabio L. Urbina, Shalini Menon, Dennis Goldfarb, Reginald Edwards, M. Ben Major, Patrick Brennwald, Stephanie L. Gupton

## Abstract

Neuronal morphogenesis involves dramatic plasma membrane expansion, likely fueled by SNARE-mediated exocytosis. Distinct fusion modes described at neuronal synapses include full-vesicle-fusion (FVF) and kiss-and-run fusion (KNR). During FVF, lumenal cargo is secreted and vesicle membrane incorporates into the plasma membrane. During KNR a transient fusion pore secretes cargo, but closes without membrane addition. In contrast, fusion modes are not described in developing neurons where plasma membrane expansion is significant. Here, we resolve individual exocytic events in developing murine cortical neurons and use new classification tools to identify four distinguishable fusion modes: two FVF-like modes that insert membrane material and two KNR-like modes that do not. Discrete fluorescence profiles suggest distinct behavior of the fusion pore with each mode. Simulations and experiments agree that FVF-like exocytosis provides sufficient membrane material for morphogenesis. We find the E3 ubiquitin ligase TRIM67 promotes FVF-like exocytosis. Our data suggest this is accomplished in part by limiting incorporation of the Qb/Qc SNARE SNAP47 into SNARE complexes and thus, SNAP47 involvement in exocytosis.

## Introduction

Membrane bound compartments define eukaryotic cells. The transfer of material between compartments requires fusion of lipid bilayers. A fusion pore is initiated by SNARE proteins assembling into a SNARE complex (**Fig 1A)**. Specific SNAREs mediate fusion between different compartments: for example, vesicle-associated membrane protein 2 (VAMP-2) and plasma membrane (PM)-associated syntaxin-1 and synaptosomal-associated protein 25 (SNAP25) facilitate docking and fusion of synaptic vesicles PM (Sudhof and Rothman, 2009; Schoch, 2001; Söllner et al., 1993). Evoked exocytosis of neurotransmitters involves multiple fusion events spatio-temporally clustered at the synapse, a diffraction-limited area. This restriction renders visualization of individual fusion events difficult. During neuronal development, however, constitutive exocytosis mediated by the same SNARE proteins is not spatially restricted and is sufficiently infrequent to resolve individual vesicle fusion events (Bello et al., 2016; Urbina et al., 2018).

**Figure 1.**
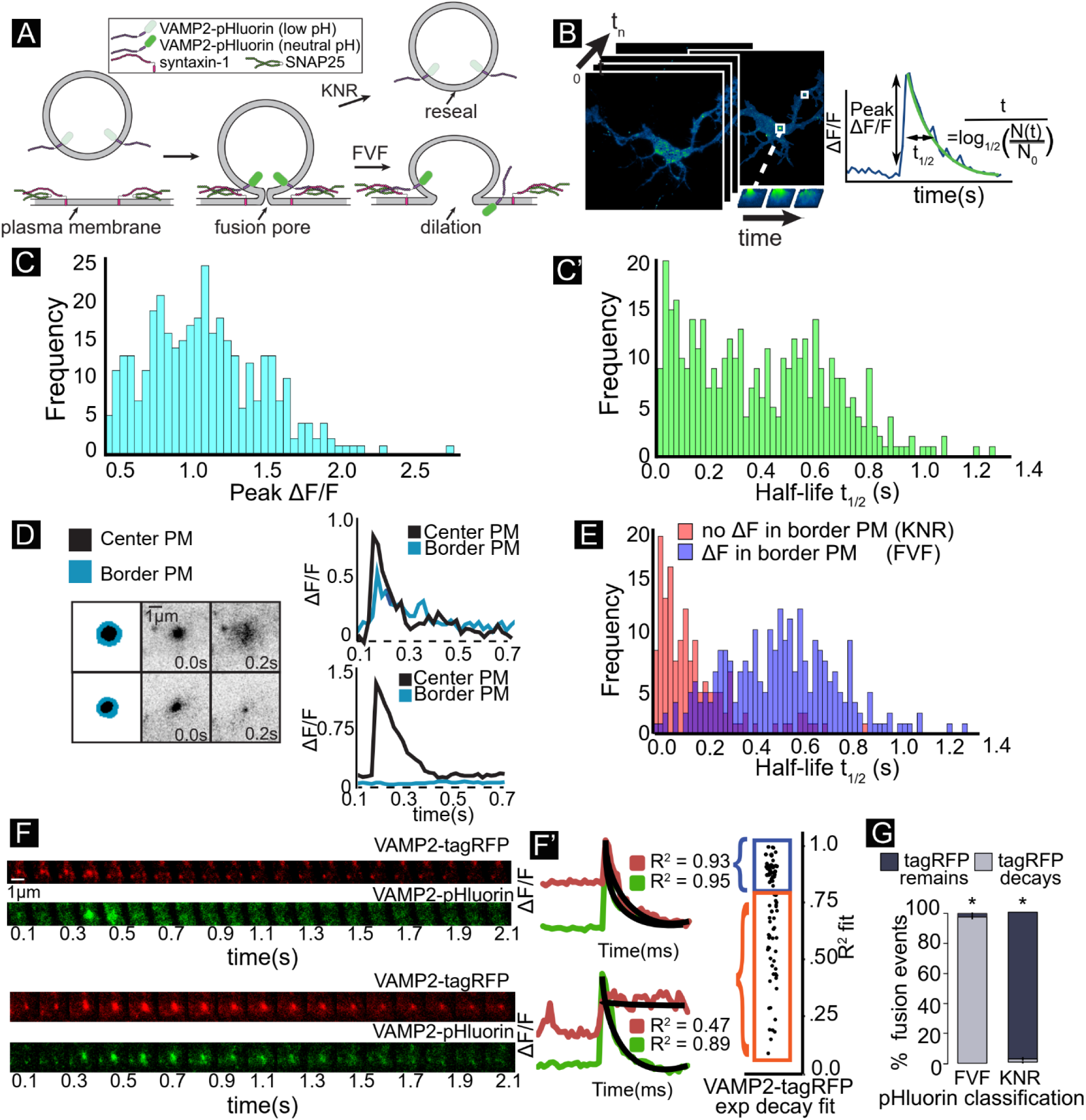
Heterogeneity in single vesicle exocytic fusion events. **A)** Schematic of exocytic fusion depicts quenching and fluorescence of VAMP2-pHluorin during full vesicle fusion (FVF) and Kiss-and-run fusion (KNR). **B)** Diagram of automated detection of exocytosis. Exocytic events (green) are detected over time. Features are extracted, including the peak ΔF/F and half-life of fluorescence. Frequency of **C)** peak ΔF/F and **C’)** fluorescence half-life (t_1/2_) of individual events. **D)** Schematic of region of exocytic event (black) and bordering plasma membrane (PM, blue). Plots of ΔF/F over time demonstrate an event with subsequent spreading of fluorescence into bordering PM (top, FVF), and an event that without spreading of fluorescence (bottom). **E)** Frequency of half-life categorized by ΔF in bordering PM. Spreading fluorescence events (ΔF, presumably FVF) have a slower fluorescence decay than those without (no ΔF, presumably KNR). **F)** Dual-color time lapse imaging of VAMP2-pHluorin and VAMP2-tagRFP reveals vesicle fusion events and vesicle fate, respectively. Example images and ΔF/F curves shows that VAMP2-tagRFP either exponentially decays after ΔF/F (top, FVF) or remains in retained vesicle (bottom, KNR). **G)** Classification of fusion events based on VAMP2-pHluorin behavior or VAMP2-tagRFP behavior agree. * represents P-value < 0.05 based on a chi-squared analysis of expected ratios if the classes were assigned randomly.

Our previous work suggested that prior to synaptogenesis, VAMP2-mediated fusion provided excess amounts of material to the expanding PM of the developing neuron, contributing to neuronal morphogenesis (Urbina et al., 2018). However, potentially not all exocytic events add membrane material and fusion mode may be a regulatable parameter of developmental exocytosis. At the synapse, two modes of exocytosis have been described, yet their contribution remains controversial (Alabi and Tsien, 2013; Albillos et al., 1997; He and Wu, 2007; Elhamdani et al., 2006). During full vesicle fusion (FVF), the fusion pore is suggested to dilate and the vesicle membrane incorporates into the PM. In contrast, during kiss and run fusion (KNR), the fusion pore opens transiently, secretes lumenal cargo, and then reseals, retaining vesicular identity (Alabi and Tsien, 2013; Albillos et al., 1997; Bowser and Khakh, 2007; Holroyd et al., 2002; Wang et al., 2003). As presumably only FVF donates membrane material, regulating the mode of exocytosis may fine tune neuronal PM expansion during development.

Fusion pore size and kinetics vary with fusion mode. How the fusion pore and thus mode of exocytosis are regulated is debated. *In vitro*, the number of SNARE complexes regulate fusion pore dilation (Bao et al., 2018; Bello et al., 2016). The composition of SNARE complexes may also regulate the fusion pore. The SNARE family contains more than 60 members (Burri and Lithgow, 2004). SNAP47 is an atypical SNAP25 family member that can substitute for SNAP25 in complexes with VAMP2 and syntaxin-1 *in vitro*. We found that an interaction between SNAP25 and the brain-enriched TRIpartite Motif (TRIM) E3 ubiquitin ligase TRIM9 prevents SNARE complex formation and consequently attenuates the rate of exocytosis and axon branching in developing neurons (Winkle et al., 2014). SNARE interacting proteins, such as TRIM9, synaptotagmins, complexins, and α-synuclein may also regulate fusion pores (Archer et al., 2002; Logan et al., 2017; Wang et al., 2003; Winkle et al., 2014). Like SNAREs, the mammalian TRIM family of ubiquitin ligases contains ∼ 70 members, with many TRIMs enriched in neurons (Napolitano and Meroni, 2012). TRIM67, a paralog of the SNAP25-interacting TRIM9, is enriched in the developing cortex and regulates axonal projections, filopodia, and spatial learning and memory (Boyer et al., 2018, 2020), but thus far has not been implicated in exocytosis. The roles that TRIM67 and SNAP47 play in VAMP2-mediated fusion in developing neurons have not been explored.

Here, we use VAMP2-pHluorin expression in developing neurons to explore modes of single exocytic events. We develop computer vision classifiers to surprisingly reveal four modes of fusion. This includes two distinct classes within both FVF and KNR, which exhibit distinct fluorescence behavior following fusion pore opening. Experimental manipulations and simulations agree that membrane provided by FVF-like modes of VAMP2-mediated exocytosis approximate PM expansion of developing neurons. The E3 ubiquitin ligase TRIM67 regulates exocytic mode by reducing SNAP47 protein and limiting SNAP47 incorporation into SNARE complexes. We show that SNAP47 co-localizes with a subset of VAMP2-mediated exocytic events and alters fusion mode.

## Results

### Heterogeneity in single vesicle fusion event kinetics

VAMP2-pHluorin labeled exocytic events on the basal surface of embryonic murine cortical neurons were imaged at two days in vitro (2 DIV) by total internal reflection fluorescence (TIRF) microscopy. VAMP2 is the most highly expressed vSNARE during neuronal development (Grassi et al, 2015; Urbina and Gupton, 2020). pHluorin is a pH-sensitive variant of GFP that is quenched in the vesicular lumen and fluorescent at extracellular pH (Miesenböck et al., 1998) (**Fig1A**). Fusion events are characterized by rapid fluorescence increase when fusion pores open (**Fig1B**) and a slower fluorescent decay. Exocytic events were automatically detected (Urbina et al., 2018). To evaluate the population of events, we examined the normalized peak change in fluorescence per event (peak ΔF/F, **Fig1B,1C**), an estimate of the relative amount of VAMP2-pHluorin per vesicle, and the event half-life (t_1/2_, **Fig1B,1C’**), which describes fluorescence decay. Unlike the peak ΔF/F, t_1/2_ exhibited a bimodal distribution, suggesting two populations of fusion events. We hypothesized these could represent two well-described modes, full-vesicle fusion (FVF) and kiss-and-run fusion (KNR, **Fig1A**)(Alabi and Tsien, 2013; Holroyd et al., 2002; Wang et al., 2003). FVF in astrocytes was shown to exhibit diffusion of VAMP2-pHluorin away from the fusion site, whereas KNR events remain fluorescent until fusion pore closure and re-acidification or retreat from the PM (Bowser and Khakh, 2007). Events with the longer t_1/2_ exhibited fluorescence spreading into the PM surrounding the exocytic event, consistent with diffusion, whereas events with a rapid t_1/2_ did not exhibit fluorescence spreading (**Fig1D**). These two responses aligned with the bimodality of t_1/2_ distribution (**Fig1E**). Thus, there is sufficient information to differentiate VAMP2-pHluorin fusion events.

Vesicles that fuse by KNR retain their identity; yet re-acidification prevents visualization of VAMP2-pHluorin. To visualize vesicles before, during, and after fusion, we imaged neurons expressing both VAMP2-pHluorin and VAMP2-tagRFP (**Fig1F**). Exponential decay of both markers confirmed diffusion of v-SNAREs during putative FVF events (**Fig1F’**, top, blue). Events without VAMP2-pHluorin diffusion maintained VAMP2-tagRFP fluorescence after fusion (**Fig1F’**, bottom, orange), indicating vesicles persisted following putative KNR events. Thus VAMP2-tagRFP fluorescence behavior agreed with pHluorin-based classification (**Fig1G**), suggesting that FVF and KNR were discernable in the pHluorin-based assay.

### Multiple unbiased classifiers converge on four exocytic modes

To explore fusion heterogeneity in an unbiased fashion, we turned to classification and clustering approaches. Most clustering algorithms rely on a single classification method. In contrast, we employed an unbiased three-pronged approach to rigorously classify vesicle fusion. Detected events were first temporally aligned to peak ΔF/F (**Fig2A, red dots**). As a ground truth method, we selected features that discern known differences between KNR and FVF based on their biological interpretation. This includes ΔF/F and t_1/2_, among other parameters (**Fig2B, see methods**). Principal Component Analysis (PCA) was performed on all features and the five principal components (PC) capturing 85% of the variance were kept (**Fig2B’**). Second, we utilized agglomerative and divisive hierarchical cluster analysis **(Fig2C)**. Hierarchical clustering summated a Euclidean distance cost (∑) between each pair of events starting at peak ΔF/F (**Fig2Ci**) and either merged the lowest-cost events iteratively (agglomerative) or iteratively divided events (divisive), creating a cluster dendrogram (**Fig2Cii**, see methods). Euclidean distance matrices were constructed to visualize the relationship and groupings of these events (**Fig2Ciii**). Finally, we utilized Dynamic Time Warping (DTW, **Fig2D**) to measure spatio-temporal dissimilarity between all pairs of ΔF/F curves. DTW allowed non-linear temporal matching between events **(Fig2Di)** to find the optimal path of smallest distance (lowest cost). DTW created a time warp matrix where each square was the Euclidean distance between respective timepoints of two ΔF/F time series (**Fig2Dii**). The optimal warp path for a pair of exocytic events was the lowest cost path through the matrix, which may be distinct from the ∑ Euclidean distance (**Fig2Dii**). The sum of the warp path, or the dynamic time warping cost, was plotted on a distance matrix, with each pixel of the distance matrix representing the sum of the warp path for a pair of exocytic events (**Fig2Diii**).

We took the plurality rules decision from a committee of the most common and best performing clustering indices (Charrad et al., 2014) to determine the number of exocytic classes or clusters (k) present in the data with confidence. Among the indices used were the gap statistic, which measured within-cluster dispersion, and the elbow method, which examined the percent variance explained as a function of the number of clusters (**Fig2E**, see methods). Unexpectedly, the committee converged on a *k* of four classes for each classification scheme instead of the two, as predicted by the literature, with eight of 20 indices selecting four classes (**Fig2F**). All four classification methods (Feature Selection, DTW, agglomerative and divisive hierarchical clustering) classified 96.3% of n = 733 exocytic events exocytic events unanimously, indicating differences between events and clusters were sufficiently robust to be discerned via multiple methods **(Fig2G)**. Inspection of events with conflicting classification revealed edge cases, with low signal-to-noise or occurrence close to or on the cell periphery (**Fig2G**, inset). One explanation for unexpected classes was grouping false-positively detected fluorescence as different from true-positive exocytic events. To control for this, we treated neurons with tetanus neurotoxin (TeNT, **Fig2H**), which cleaves VAMP2 and blocks VAMP2-mediated fusion (Link et al., 1992), including VAMP2-pHluorin mediated events (Urbina et al, 2018). All four classes were sensitive to TeNT, indicating that all four classes contain bona fide exocytic events.

**Figure 2:**
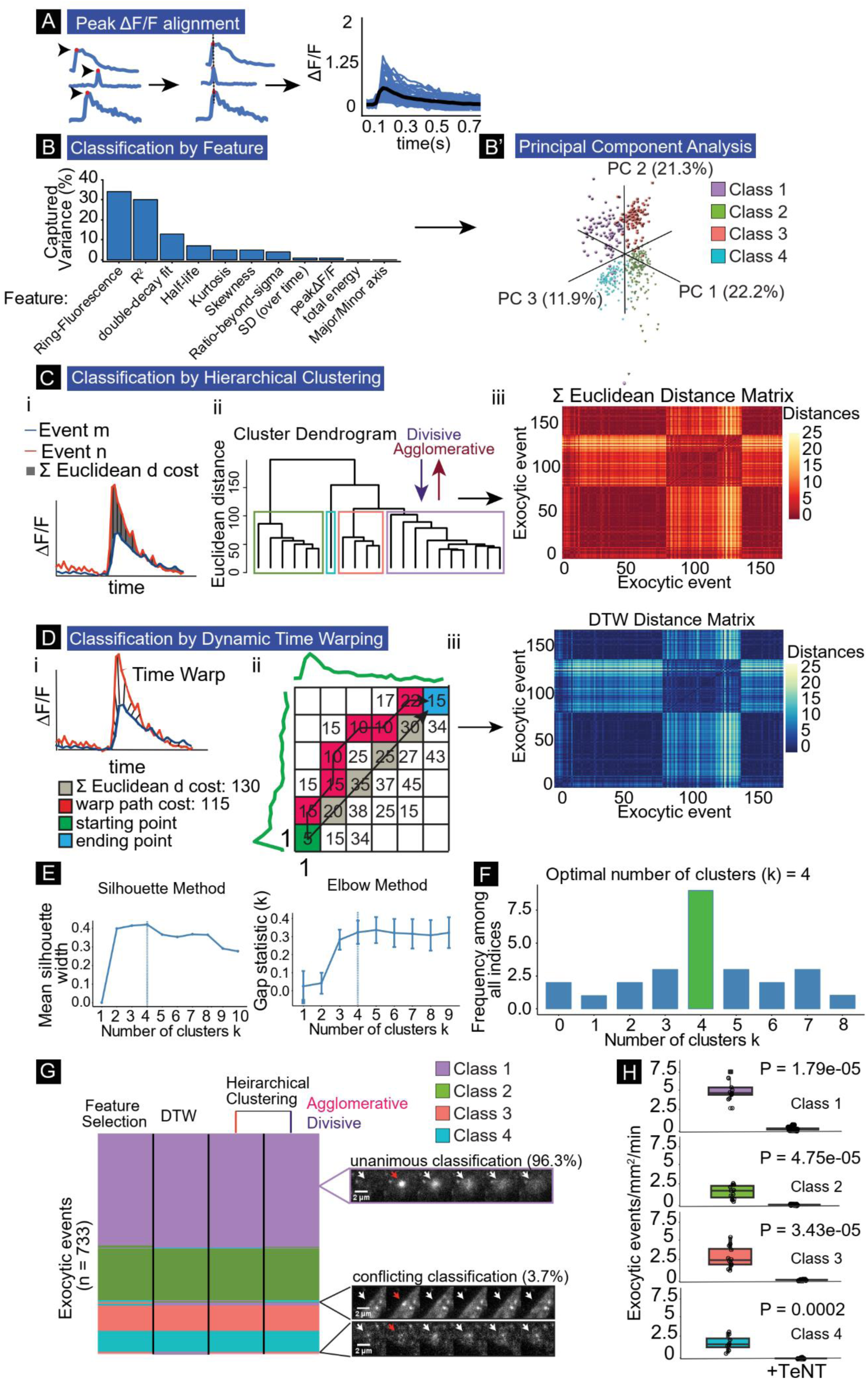
Multiple unbiased classifiers converge on four exocytic modes. **A)** ΔF/F curves are temporally registered to peak ΔF/F (red dot). Registered ΔF/F plot showing average (black) and individual events (blue). **B)** Features that represent % of captured variance from PCA. **B’)** Three principal components plotted revealing four exocytic classes. **C)** Event classification by hierarchical clustering. i) ΔF/F plot depicting the summed (∑) Euclidean distance between two exocytic events, m and n. ii) Dendrogram of ∑ Euclidean distances between multiple events, linked by agglomerative or divisive hierarchical clustering revealing four exocytic classes. iii) Each pixel in distance matrix is the ∑ Euclidean distance between two ΔF/F curves (n=733). **D)** Event classification by Dynamic Time Warping (DTW). i) ΔF/F plot depicting time warped distance between two exocytic events, m and n. ii) A matrix of Euclidean distances measured at all time point comparing a pair of exocytic events, starting at peak ΔF/F (green square, co-ordinates 1,1). Unlike the ∑ Euclidean distance (gray), DTW uses the warp path with the lowest cost (red squares) to reach the ending point (blue square). The sum of the total warp path is the DTW cost per pair of exocytic events. iii) Each pixel in distance matrix is the DTW cost between two ΔF/F curves (n = 733). **E)** Two example indices, silhouette and elbow methods, used in the plurality rules committee suggest four exocytic classes. **F)** Decision histogram of the committee of indices, with the plurality of indices choosing four exocytic classes (8/20). **G)** Classification comparison of each of the methods of clustering. 96.3% of 733 events were classified unanimously. Purple inset: representative unanimously classified exocytic event. Black inset: representative events with conflicting classification. Cell edge effects (top black inset) or low signal-to-noise ratio (bottom black inset) account for the majority of conflict. White arrows denote exocytic events. Red arrows denote peak ΔF/F. **H)** Frequency of all event classes drops to near zero with tetanus toxin (TeNT) treatment (Welch’s t-test, n = 12 cells per experiment).

### Distinguishing features of four modes of exocytosis

We next characterized each exocytic class (**Fig3**). Class averages and representative image sequences revealed distinct fluorescence profiles (**Fig3A, Movie 1**). Class 1 and 2 exhibited an instantaneous fluorescence decay after peak ΔF/F (fusion pore opening). In contrast, class 3 and 4 exhibited a delay between peak ΔF/F and the onset of fluorescent decay, observed as a plateau in ΔF/F prior to decay. A single exponential decay fit well to the fluorescence decay in classes 1 and 2 (**Fig3B**), but was a poor fit for classes 3 and 4 (**Fig3C**). Fitting sequential exponentials resulted in a higher R^2^ (**Fig3D**). t_1/2_ for class 3 and 4 were calculated from the second exponential, which revealed no differences in t_1/2_ between class 1 and 3 or class 2 and 4 (**Fig3E**).

Fluorescence decay following FVF is due to VAMP2-pHluorin diffusion into the bordering PM (**Fig3F**). The PM surrounding class 1 and 3 events exhibited a fluorescent peak followed by an exponential decay with the same t_1/2_ as the center pixels **(Fig3G,H, Movie 1)**, consistent with diffusion and FVF. In contrast, VAMP2-pHluorin fluorescence did not increase in the PM surrounding class 2 and 4 events (**Fig3F,G**), suggesting these may be KNR. Loss of fluorescence following KNR is due to either vesicle-reacidification or vesicle retreat from the evanescent wave. The t_1/2_ of class 2 and 4 did not change after increasing the evanescent wave penetration (**FigS1A**), indicating significant fluorescence loss was not due to vesicle retreat from the evanescent wave. In contrast, pH buffering the media with cell-impermeable HEPES slowed fluorescence decay of class 2 and 4 events, but not class 1 and 3 (**Fig3E**). This suggested that HEPES entered class 2 and 4 vesicles before resealing and retarded reacidification, consistent with KNR. The t_1/2_ of HEPES-sensitive events were similar to single-vesicle KNR events measured in astrocytes (Bowser and Khakh, 2007), but faster than reported at synapses (Pradeep and Ryan, 2006). These results suggest class 2 and 4 are KNR and that reacidification rates may differ in developing and mature neurons. Based on the mechanisms of fluorescence decay and either the instantaneous (i) or delayed (d) onset of fluorescence decay after peak ΔF/F, we named the classes FVFi, (class 1), KNRi (class 2), FVFd (class 3), and KNRd (class 4).

**Figure 3:**
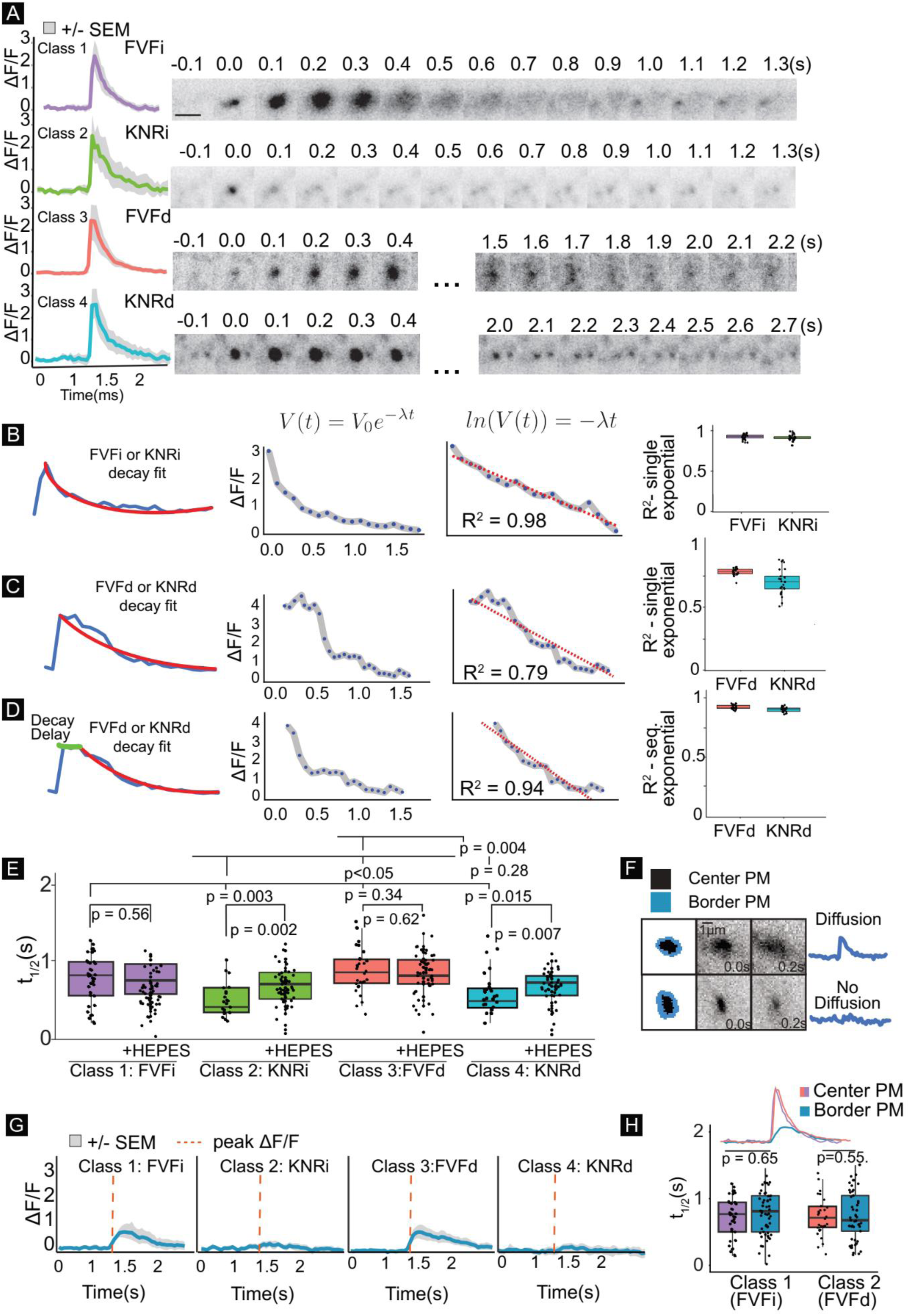
Distinguishing features of four modes of exocytosis. **A)** Mean fluorescent curves +/- SEM (gray) and representative images from each class (right). **B-C)** Example single exponential decay fit (V(t) = V_0_e^- λt^) to a class 1 or 2 instantaneous (i) events (B) or to class 3 or 4 delay (d) events (C). The log of an exponential decay curve (ln(V(t))=- λt) is linear, and the linear regression fit (R^2^) is an indicator of how good of a fit the exponential decay curve is (center graph). Box plots (right) of R^2^ of class 1 (FVFi) and class 2 (KNRi) events show a good fit (B) whereas fit to class 3 (FVFd) and class 4 (KNRd) are poor. **D)** Sequential exponential decay curves better fit to FVFd and KNRd. **E)** t_1/2_ of each class untreated or treated with HEPES. Class 2 and 4 are HEPES sensitive, indicating they represent KNR-like events (n = 14 cells per condition; Welch’s t-test followed by Benjamini-Hochberg correction). **F)** Schematic of region of exocytic event (black) and bordering PM (blue). Plots of border PM ΔF/F over time demonstrate an event with fluorescence spreading (top, FVF), and an event without (bottom). **G)** Class means +/- SEM of ΔF/F in PM bordering exocytic events. Red dotted line indicates peak ΔF/F of center of event. Red dotted line indicates peak ΔF/F of the center of event. **H)** Half-life of fluorescence decay (t_1/2_) in PM bordering FVFi or FVFd events (border pixels) is not different from the t_1/2_ of the exocytic event (center pixels) (n = 14 cells per condition; Welch’s t-test).

Surface levels of VAMP2-pHluorin varied between cells and between experiments, but did not alter the distribution exocytic modes (**FigS1B**,**C**). The consistency in the distribution of modes regardless of VAMP2 expression level or biological replicate suggested that cellular VAMP2 levels did not alter exocytic modes. There was a stratification of peak ΔF/F between modes (**FigS1D**). FVFi and KNRi events had a higher peak ΔF/F than FVFd and KNRd, suggesting vesicular VAMP2 levels influence fusion mode.

To further ensure the four exocytic modes were not an artifact of VAMP2pHluorin, we employed alternative fusion reporters. Transferrin receptor (TfR) attached to a pH-sensitive red fluorophore pHuji showed 100% overlap with VAMP2-pHluorin fusion events **(FigS2A)**. We thus detected and analyzed exocytic events based of TfR-pHuji fluorescence and diffusion behavior (see Methods). Using all three classifiers with the committee of indices, four classes of exocytosis were detected for TfR-pHuji with identical ratios to VAMP2-pHluorin (**Fig S2B,C**). The t_1/2_ of KNRi and KNRd for TfR-pHuji was the same as VAMP2-pHluorin (**FigS2D,E**), consistent with the similar pH sensitivity of the fluorophores (Martineau et al, 2017), supporting that decay was due to reacidification. In contrast, the t_1/2_ of FVFi and FVFd were different between the two exocytic markers (**FigS2D,E)**, which can be explained by the distinct size and diffusion behaviors of VAMP2 and TfR (Di Rienzo et al., 2013; Fujiwara et al., 2002, 2016; Jaqaman and Grinstein, 2012; Lenne et al., 2006). This further supports diffusion-dependent fluorescence decay.

As an additional marker of exocytosis, we turned to the vSNARE VAMP7, which mediates exocytosis of a discrete population of vesicles in developing neurons, albeit at lower frequencies than VAMP2 (Gupton and Gertler, 2010; Urbina et al.,2018). VAMP7-mediated fusion is autoinhibited by the longin doman of VAMP7 (Burgo et al., 2013; Martinez-Arca et al. 2001); a VAMP7-pHluorin construct lacking the longin domain (VAMP7ΔLD) increased the frequency of exocytosis (Urbina et al, 2018) sufficiently to identify exocytic modes. Four distinct modes of VAMP7ΔLD exocytosis were detected by the three classifiers for this discrete vesicle population. Interestingly the relative abundance of each fusion mode was distinct from VAMP2-pHuorin (**FigS2F**). Together the VAMP7ΔLD and TfR data confirmed the existence of four exocytic modes.

### Expression of a truncated VAMP2 alters exocytic mode

The VAMP2 transmembrane domain catalyzes fusion initiation and pore expansion (Dhara et al., 2016). Capacitance measurements demonstrated that deletion of the transmembrane domain stabilized narrow fusion pores and blocked full fusion (Guček et al., 2016). To determine if fusion pore behavior was distinguishing exocytic mode, we expressed tagRFP-VAMP2 lacking the transmembrane domain (VAMP2^1-96^, **FigS3A**) along with VAMP2-pHluorin. VAMP2^1-96^ reduced the frequency of exocytic events (**FigS3B, Movie 2**). Intriguingly, VAMP2^1-96^ specifically reduced the frequency of FVFi and FVFd events (**FigS3C**). This shifted the distribution of exocytic events toward KNRd and KNRi (**FigS3D**), aligning with capacitance measurements (Guček et al., 2016). VAMP2^1-96^ specifically decreased the frequency of FVFi and FVFd and not KNRi and KNRd, suggesting that the FVF-like classes may be mechanistically distinct from KNR.

### TRIM67 biases exocytic mode towards full-vesicle fusion

Experiments with TeNT (**Fig2H**), HEPES (**Fig3E**), VAMP2-tagRFP (**Fig1F**), VAMP2^1-96^ (**FigS3**), VAMP7ΔLD-pHluorin **(FigS2F)**, and TfR-pHuji **(FigS2B-E)** support the conclusion that each exocytic class harbors bona fide and distinct exocytic events, yet the biological relevance of and molecular mechanisms influencing exocytic modes are unclear. Our previous work identified the E3 ubiquitin ligase TRIM9 as a novel regulator of neuronal exocytosis (Urbina et al., 2018; Winkle et al., 2014). Although deletion of *Trim9* elevated the frequency of exocytosis (Urbina et al., 2018; Winkle et al., 2014), there was not a change in the ratio of classes **(Fig4A, Movie 3)**. TRIM67 is a TRIM9 paralog enriched in developing neurons (Boyer et al., 2018). Unlike *Trim9*, deletion of *Trim67* altered the mode but not the frequency of exocytosis; KNRi and KNRd increased two-fold, whereas FVFi and FVFd decreased (**Fig4A,B**). The distribution of exocytic classes within the soma was not distinct from the whole neuron in both *Trim67*^*-/-*^ and *Trim67*^*+/+*^ neurons, however FVFd events were enriched in neurites of both **(Fig4C)**. These data indicate that exocytic frequency and mode are independently regulated, in this case by distinct E3 ligases.

During morphogenesis neuronal surface area increases (**Fig4D**). Our previous experimental and modeling work suggested that VAMP2-mediated exocytosis provided excess material for PM expansion, which was partially balanced by clathrin-mediated endocytosis (Urbina et al., 2018) (**Fig4**). The model used empirically measured surface areas of neurons, non-coated, and clathrin-coated vesicles, along with the frequencies of exocytosis and endocytosis to predict plasma membrane expansion. A caveat was the assumption that all exocytic events supplied PM material. Upon considering that KNRi and KNRd comprise ∼22% of events (**Fig4A**), we updated the model with exocytic modes (**Fig4F**) and new starting surface area measurements. This demonstrated that accounting for KNRi and KNRd improved the similarity of predictions to measured values of surface area (**Fig4H vs 4E**). We extended the simulation to 3 DIV (**Fig4I**). The modeled neurons matched the measured surface area well, however a population of neurons with larger surface areas arose at 3 DIV that were not reflected in simulations.

**Figure 4:**
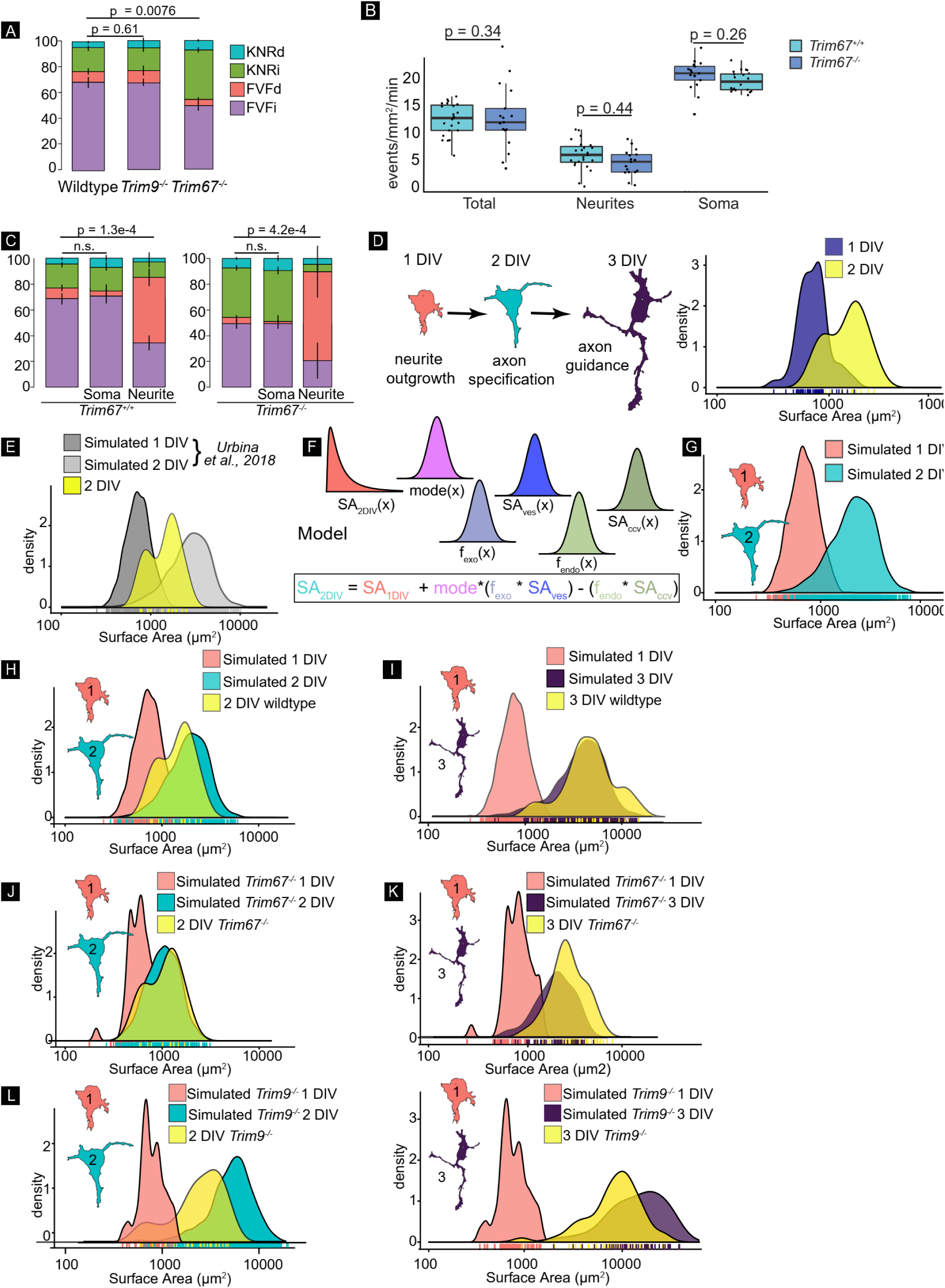
TRIM67 biases exocytic mode towards full-vesicle fusion. **A)** Proportion of KNRd, KNRi, FVFd, and FVFi in wildtype, *Trim9*^*-/-*^, or *Trim67*^*-/-*^ neurons (n = 15 neurons per condition; n = 4 biological replicates; multivariate linear regression). **B)** Frequency of exocytic events in the whole neuron, soma, and neurites of *Trim67*^*+/+*^ or *Trim67*^*-/-*^ neurons (n = 15 neurons per condition; n = 3 biological replicates; Welch’s t-test followed by Benjamini Hochberg correction). **C)** Proportion of KNRd, KNRi, FVFd, and FVFi VAMP2-pHluorin exocytic events in the whole neuron, soma, and neurites of *Trim67*^*+/+*^ or *Trim67*^*-/-*^ neurons (n = 15 neurons per condition; multivariate linear regression). **D)** Schematic of neuronal morphogenesis in vitro (left) and empirically measured surface areas from GFP-CAAX expressing wildtype neurons at indicated time points (right). Surface area calculations were performed as described in (Urbina et al., 2018). **E)** Simulated 1 and 2 DIV neurons using the model from (Urbina et al., 2018) that considers all exocytic events FVF. Empirically measured surface area of cortical neurons *in vitro* plotted in yellow. **F)** Overview of new cell model that includes the mode of exocytosis for simulating cell growth. Surface area at 2 DIV (SA_2DIV_) was estimated by first generating a population of neurons at 1 DIV (SA_1DIV_) based on empirical measurements of neurons at 1DIV. The distribution of modes (mode), frequency of exocytosis (f_exo_), surface area of vesicles (SA_ves_), frequency of endocytosis (f_endo_), and surface area of clathrin-coated vesicles (SA_ccv_) were incorporated into estimating the surface area expansion. **G)** Simulated surface areas at 1 and 2 DIV. Surface areas at 1 DIV are simulated based on empirical measurements of surface area at that time point. Surface areas at 2 DIV use the 1 DIV simulation and model described in F. **H)** Simulated surface areas from G, with empirically measured surface areas at 2 DIV overlaid in yellow demonstrate a better fit that the model that only considers FVF in E. **I)** Simulated neurons at 3 DIV (purple) with measured surface area at neurons 3 DIV overlaid in yellow. **J)** Predicted PM growth simulated over 1 DIV using the relative proportion of modes from *Trim67*^*-/-*^ neurons with empirically measured *Trim67*^*-/-*^ neurons overlaid in yellow. **K)** PM growth of *Trim67*^*-/-*^ neurons simulated at 3 DIV, compared to empirically measured neurons (yellow overlay). **L)** Simulations of PM growth at 2 and 3 DIV using the distribution of frequencies from *Trim9*^*-/-*^ cortical neurons as compared to empirically measured *Trim9*^*-/-*^ neurons (yellow overlay).

Since deletion of *Trim67* altered the distribution of exocytic mode, we simulated growth of *Trim67*^*- /-*^ neurons (**Fig4J,K**). Incorporation of increased KNR and less FVF predicted a reduction in surface area expansion in *Trim67*^*-/-*^ neurons at both 2 DIV and 3 DIV. Experimentally measured surface area of *Trim67*^*-/-*^ neurons at 2 and 3 DIV agreed **(Fig4J,K)**. This suggested a bias in exocytic mode was sufficient to alter surface area. Consistent with decelerated morphogenesis, callosal axons in *Trim67*^*-/-*^ mice were delayed in midline crossing (Boyer et al, 2020). Simulations using the increased exocytic frequency of *Trim9*^*-/-*^ neurons predicted increased neuronal surfaces areas, which was confirmed empirically at 2 and 3 DIV (**Fig4L**). *Trim9*^*-/-*^ simulations did not fit as well, overestimating the increase in neuron size, suggesting compensatory mechanisms exist which were not captured in our model. Despite this, the remarkable fit between simulated and empirically measured data suggested that we have captured relevant factors responsible for developmental neuronal growth.

### The t-SNARE SNAP47 interacts with TRIM67 and localizes to VAMP2-mediated exocytic events

Time-lapse TIRF microscopy revealed that TRIM67 did not localize to VAMP2-mediated exocytic events (**Fig5A; Movie 4**). The distance between VAMP2-pHluorin exocytic events and TRIM67 were compared to theoretically randomly distributed points using Ripley’s L analysis, which indicated they were below the threshold for colocalization (red dotted line). Expression of RFP-tagged TRIM67 in *Trim67*^*-/-*^ neurons **(FigS4A)** rescued the ratio of exocytic modes (**FigS4B**) without changing the frequency of exocytosis (**FigS4C**). These data confirmed that TRIM67 biased the mode of exocytosis toward FVFi and FVFd and away from KNRi and KNRd. However, the lack of colocalization between TRIM67 and VAMP2-pHluorin exocytosis suggested that TRIM67 likely biased exocytic mode remotely.

Because few interaction partners of TRIM67 are known, we used the BioID method (Roux et al., 2012) to identify candidate TRIM67 interacting partners that regulate exocytosis. One interesting candidate and the only SNARE protein significantly enriched (∼4 fold) over the negative control (Myc-BirA*, **Table S1**) was SNAP47, a t-SNARE in the SNAP25 family. SNAP47 forms thermally stable SNARE complexes and competes with SNAP25 in SNARE complex formation in vitro (Holt et al., 2006). Although SNAP47-containing SNARE complexes are capable of fusion, they are significantly less efficient in liposomal fusion assays (Holt et al., 2006). Immunoprecipitation of Myc-SNAP47 co-precipitated GFP-TRIM67, but not GFP-TRIM9 from HEK293 cells (**Fig5B**), confirming that TRIM67 and SNAP47 interact. Live TIRF microscopy revealed that TRIM67-GFP puncta colocalize with SNAP47-tagRFP, particularly at the periphery of neurons (**Fig5C**, arrows**; Movie 5**). Time-lapse TIRF microscopy also revealed that SNAP47 co-localized with a subset (28%) of VAMP2-pHluorin exocytic events (**Fig5D; Movie 6**). Although we did not perform three color imaging, these data suggested that SNAP47 and TRIM67 interacted at sites distal from exocytic events.

**Figure 5:**
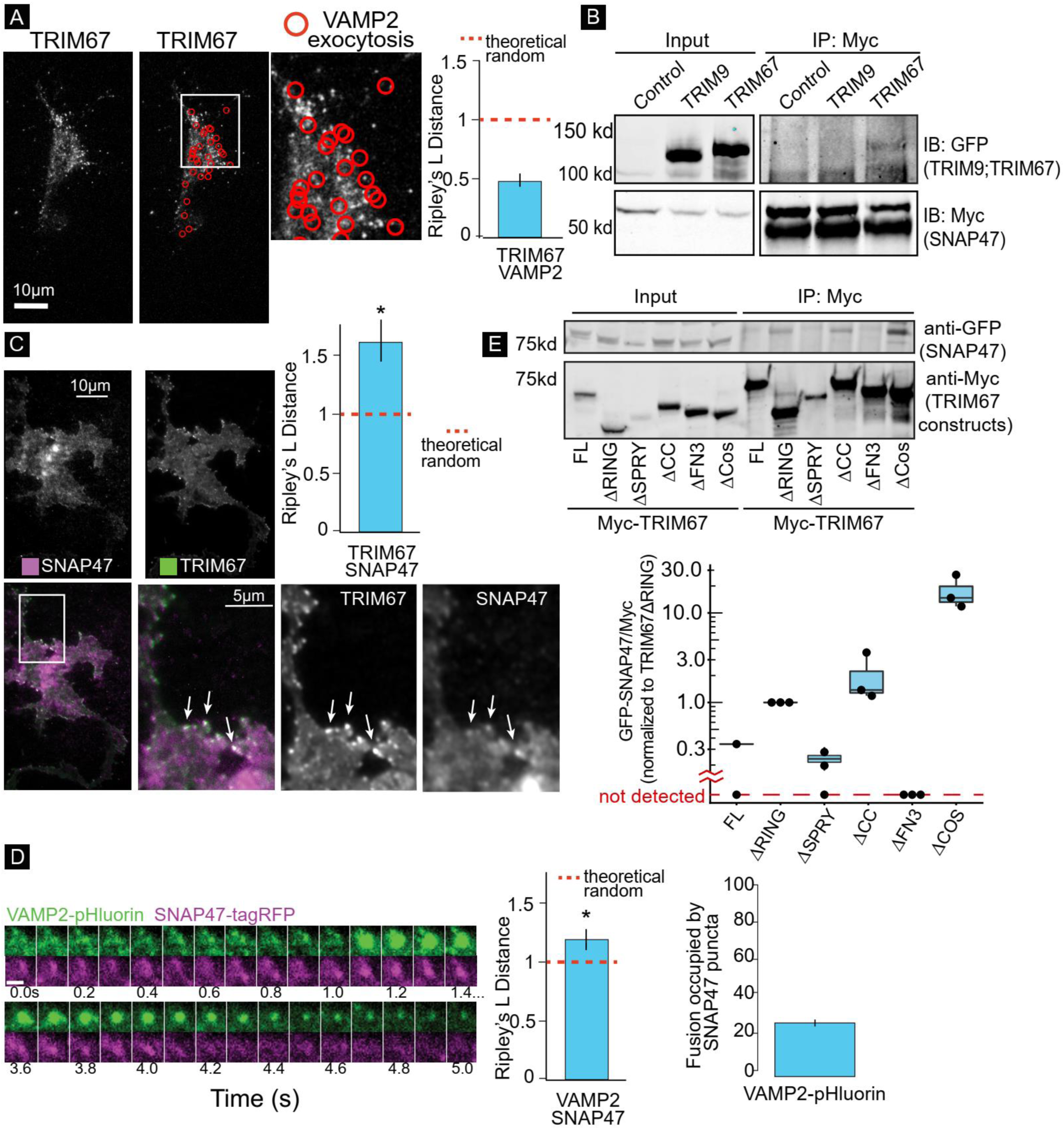
The t-SNARE SNAP47 interacts with TRIM67 and localizes to VAMP2-mediated exocytic events. **A)** Representative images and Ripley’s L Distance co-localization analysis of TRIM67-tagRFP and VAMP2-pHluorin events (red circles) n = 15 cells; n = 4 biological replicates; value lower than theoretical random suggest not significantly colocalized. **B)** Immunoprecipitation (IP:Myc) of Myc-SNAP47 from HEK293 cells expressing Myc-SNAP47 and either GFP-TRIM67 or GFP-TRIM9, blotted for GFP and Myc. GFP-TRIM67 co-IPs with Myc-SNAP47, whereas GFP-TRIM9 was not detectable in the Myc-SNAP47 IP. **C)** Average of 10 frames of images and co-localization analysis of SNAP47-tagRFP and TRIM67-GFP in cortical neurons at 2 DIV (n = 14 cells; n = 3 biological replicates; value above theoretical line are significantly colocalized (*); analysis described in methods). **D)** Example of VAMP2-pHluorin and SNAP47-tagRFP co-localization during an exocytic event. n = 16 cells; value above theoretical line are significantly colocalized (*). Scale bar = 1µm. **E)** Structure-function co-IP assays. MycTRIM67 construct IP blotted for GFP-SNAP47 and Myc. Graph represents relative GFP-SNAP47/Myc-TRIM67 constructs normalized to Myc-TRIM67ΔRING; blots without a detectable band of GFP-SNAP47 represented below graph break.

### Multiple domains of TRIM67 modulate the interaction with SNAP47 and exocytic mode

These data suggested that SNAP47 was a promising candidate to be the intermediary between TRIM67 and the fusion apparatus in regulating exocytic mode. To determine domains of TRIM67 important for interacting with SNAP47, we performed co-immunoprecipitation assays using TRIM67 domain mutants expressed in *TRIM67*^-/-^ HEK293 cells (Boyer et al., 2018)(**Fig5E**). Similar structure:function experiments were performed in *Trim67*^*-/-*^ neurons to examine exocytic mode using RFP-TRIM67 domain mutants (**FigS4B,C**). MycTRIM67ΔFN3 failed to precipitate detectable levels of GFP-SNAP47 (**Fig5E**) or rescue the ratio of exocytic modes (**FigS4B**). MycTRIM67ΔSPRY precipitated similar amounts of GFP-SNAP47 compared to TRIM67 and rescued exocytic mode (**Fig5E**,**S4B**). In contrast, more SNAP47 was detected in constructs lacking the ligase containing RING domain (MycTRIM67ΔRING) or multimerization coiled-coil domain (MycTRIM67ΔCC), yet these mutants failed to rescue the ratio of exocytic modes. Similarly, mutation of conserved Zn-coordinating cysteines required for ligase activity (TRIM67LD,(Boyer et al., 2020)) failed to rescue exocytic mode (**FigS4B**). None of these mutants altered the frequency of exocytosis (**FigS4C**). Deletion of the COS domain (TRIM67ΔCOS) however exhibited a unique phenotype; it decreased the frequency of exocytosis but did not rescue the ratio of exocytic modes (**FigS4B,C**). Interestingly, MycTRIM67ΔCOS precipitated much more GFP-SNAP47 (**Fig5E**). These experiments were consistent with the hypothesis that a transient interaction between TRIM67 and SNAP47 was required for appropriate regulation of exocytic mode.

### TRIM67 alters exocytosis via modulated SNAP47 protein levels

Analysis of cell lysates revealed a 1.5x fold increase in SNAP47 protein in *Trim67*^*-/-*^ neurons (**Fig6A**), suggesting that TRIM67 regulated SNAP47 expression or stability. A cycloheximide chase revealed that SNAP47 was degraded at the same rate in *Trim67*^*+/+*^ and *Trim67*^*-/-*^ cortical neurons (**FigS5A**). Bortezomib or chloroquine treatments indicated SNAP47 degradation was proteasome-dependent and lysosome independent (**FigS5A**) in both *Trim67*^*+/+*^ and *Trim67*^*-/-*^ cortical neurons. To assess SNAP47 ubiquitination, we performed denaturing immunoprecipitation of GFP-SNAP47 in *TRIM67*^*-/-*^ HEK293 cells co-expressing HA-ubiquitin and either Myc or Myc-TRIM67 **(FigS5B)**. The ratio of SNAP47-Ub:SNAP47 was not altered by loss of TRIM67. Altogether these data indicated that SNAP47 ubiquitination and degradation were not detectably altered by loss of TRIM67. Currently the precise mechanism by which loss of TRIM67 results in elevated SNAP47 protein is unknown. However, we hypothesized that the elevated SNAP47 in *Trim67*^*-/-*^ neurons may alter the ratio of exocytic modes. Like deletion of *Trim67* (**Fig4B**), overexpression of SNAP47 in *Trim67*^*+/+*^ neurons did not alter the frequency of exocytosis (**Fig6B**). However, overexpression of SNAP47 increased KNRd and decreased FVFi and FVFd (**Fig6C**). Strikingly, the delay time before onset of fluorescent decay increased for KNRd and FVFd (**Fig6D**), which was specifically pronounced in events where SNAP47-tagRFP was detectable (red data points). Once fluorescence decay initiated, overexpression of SNAP47 did not affect half-life (**Fig6E**). These findings indicated increased SNAP47 promoted KNRd and was sufficient to shift exocytic mode. To determine if increased SNAP47 was required to shift exocytic mode in *Trim67*^*-/-*^ neurons, we reduced SNAP47 protein levels using siRNA (**Fig6F,F’**). siRNA resulted in ∼80% knockdown of SNAP47 (**Fig6F,F’**). Knockdown of SNAP47 did not alter the frequency of exocytosis (**Fig6G**), but decreased KNRd and KNRi and increased FVFi and FVFd in *Trim67*^*-/-*^ neurons (**Fig6H**). The significant differences in exocytic modes observed with both increased and decreased SNAP47 suggested that SNAP47 regulated the mode of exocytosis.

**Figure 6:**
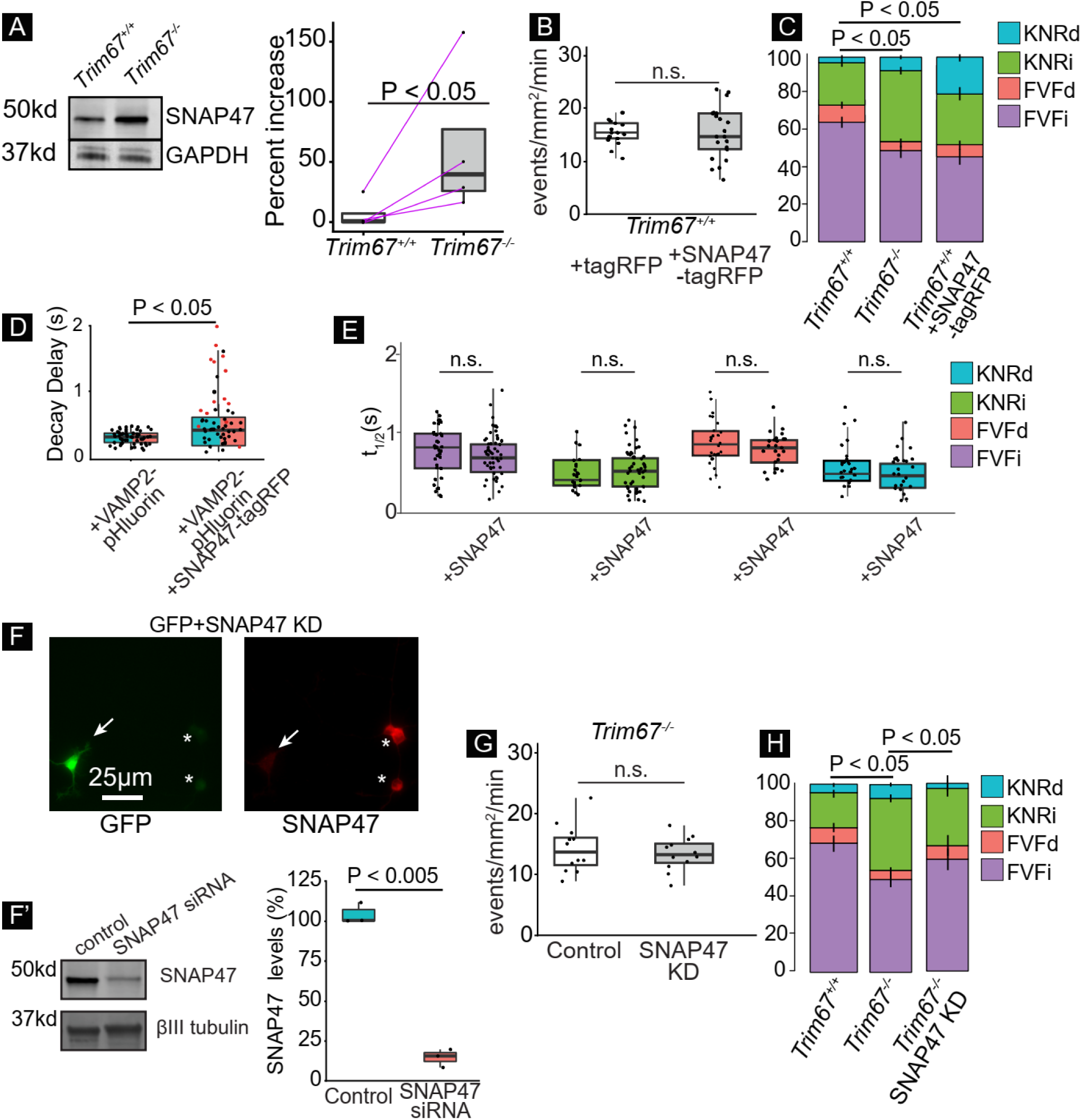
TRIM67 alters exocytosis by modulating SNAP47 protein levels. **A)** Immunoblot of SNAP47 and GAPDH from lysate of *Trim67*^*+/+*^ and *Trim67*^*-/-*^ cortical neurons at 2 DIV. Quantification of percent increase of SNAP47/GAPDH quantified from 4 blots (right; log paired-t-test, analysis in methods). **B)** Frequency of VAMP2-pHluorin exocytosis +/- tagRFP or SNAP47-tagRFP is not significantly differently (n = 16 neurons per condition, n = 3 biological experiments, Welch’s t-test). **C)** Relative proportions of exocytic modes in *Trim67*^*+/+*^ neurons, *Trim67*^*-/-*^ neurons, or *Trim67*^*+/+*^ neurons expressing SNAP47-tagRFP (n = 16 neurons per condition; n = 3 biological replicates; multivariate linear regression). **D)** Quantification of time after peak ΔF/F before fluorescence decay onset of KNRd and FVFd events (n = 17 neurons; n = 4 biological replicates; Welch’s t-test). **E)** t_1/2_ of each exocytic class +/- SNAP47-tagRFP was not different (n = 17 neurons per condition; Welch’s t-test with Benjamini-Hochberg correction). **F)** Example SNAP47 immunocytochemistry images in neurons expressing siRNA for SNAP47 knockdown (KD) (GFP+, arrow), or without (*). Cells were transfected with an eGFP expression plasmid and SNAP47 siRNA followed by immunostaining of SNAP47; GFP positive neurons did not stain for SNAP47, suggesting knockdown of SNAP47. **F’)** Immunoblot and quantification of SNAP47 and βIII tubulin protein levels in SNAP47 knockdown compared to control (n = 3 experiments; Welch’s t-test). **G)** Frequency of VAMP2-pHluorin exocytosis +/- SNAP47 siRNA is not different (n = 12 neurons per condition; Welch’s t-test). **H)** Relative proportions of each exocytic class in *Trim67*^*+/+*^ neurons, *Trim67*^*-/-*^ neurons, or *Trim67*^*-/-*^ neurons with SNAP47 knockdown (n = 12 neurons per condition, n = 3 biological replicates; multivariate linear regression).

### SNAP47 forms more SNARE complexes in *Trim67*^*-/-*^ neurons

The increased SNAP47 protein in *Trim67*^*-/-*^ neurons altered the ratio of SNAP47 to SNAP25 protein **(Fig7A)**. Endogenous immunoprecipitation of SNAP47 from membrane fractions of *Trim67*^+/+^ and *Trim67*^-/-^ neurons showed that SNAP47 interacted with syntaxin-1 and VAMP2 (**Fig7B**). We thus hypothesized that SNAP47 altered exocytosis by forming more SNARE complexes in *Trim67*^*-/-*^ neurons. To test this, we compared SDS-resistant SNARE complexes, which run at higher molecular weights than their monomeric constituents in both *Trim67*^+/+^ and *Trim67*^-/-^ neurons. To reduce post-fusion SNARE disassembly and enhance detection, neurons were treated with NEM. This was compared to both NEM/DTT treatment at 37°C or incubation at 100 °C, which allow or promote SNARE disassembly, respectively (Banerjee et al., 1996). VAMP2, SNAP25, and syntaxin-1 exhibited similar proportions of complex:monomer between genotypes. In contrast, *Trim67*^*-/-*^ neurons exhibited increased amounts of complex:monomer in *Trim67*^*-/-*^ neurons (**Fig7C**). This suggested that TRIM67 actively inhibited the incorporation of SNAP47 into SNARE complexes and thus, SNAP47-mediated fusion events.

**Figure 7:**
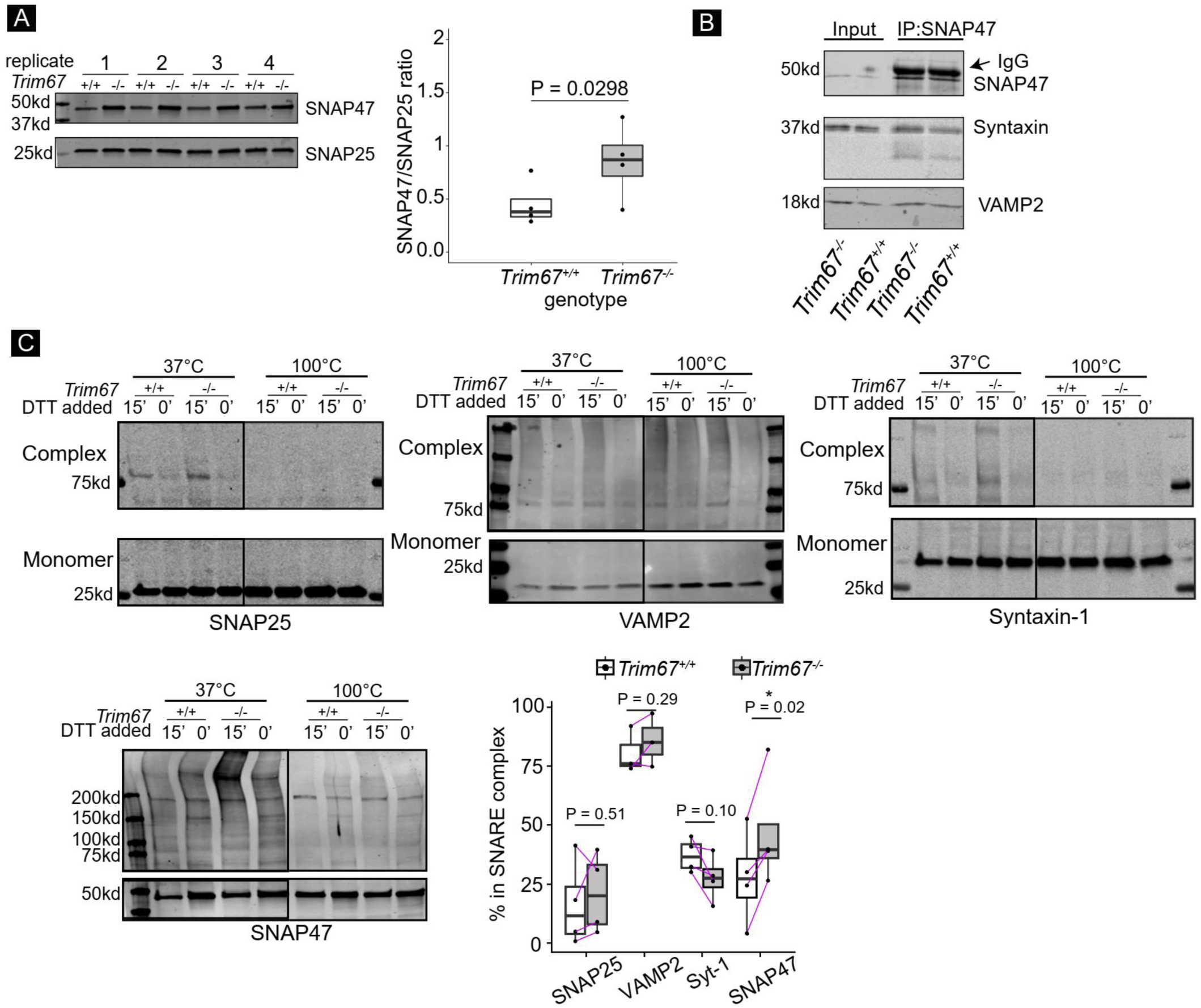
TRIM67 reduces SNAP47 incorporation into SNARE complexes. **A)** Immunoblot and quantification of SNAP47 and SNAP25 from *Trim67*^*+/+*^ and *Trim67*^*-/-*^ neuronal lysates at 2 DIV (four biological replicates). Quantified is the ratio of SNAP47/SNAP25 intensity, log-ratio paired t-test. **B)** IP of endogenous SNAP47 in neuronal membrane fractions at 2 DIV immunoblotted for SNAP47, syntaxin-1, and VAMP2. **C)** Immunoblot of monomeric and high molecular weight SNARE proteins, interpreted as SNARE complexes. Cortical neurons were treated with NEM (15 minutes) to preserve SNARE complexes followed by DTT (15 minutes) to quench NEM, or NEM+DTT as a negative control (30 minutes). Increased intensity of bands above 37kD (50kD for SNAP47) in the presence of NEM-only at 37°C suggest formed SNARE complexes. Quantification of % of each SNARE protein in complex (intensity of lane above 37kD or 50kD/(monomer + intensity of lane); n = 4 blots, log-ratio paired t-test; see methods).

## Discussion

Here we developed an unsupervised set of classifiers based on VAMP2-pHluorin fluorescence to identify four distinct exocytic classes in developing cortical neurons. These four classes were independent of cellular VAMP2 levels and detected with other markers and vesicle types. Distinguishable mechanisms of VAMP2-pHluorin fluorescence decay that were either pH-sensitive or diffusion-dependent suggested that two classes were KNR-like and two classes were FVF-like. The delayed or instantaneous onset of fluorescence decay after fusion pore opening further distinguished exocytic classes into KNRd, KNRi, FVFd, and FVFi. Even though the diffraction-limited resolution of our microscope obfuscates visualization of fusion pore properties other than opening, the alignment of our experiments using VAMP2^1-96^ with capacitance measures supports the hypothesis that the visualized delay in fluorescence decay is either a delay in fusion pore dilation (FVFd) or delay in fusion pore closure (KNRd). Deletion of *Trim67* increased KNRi and KNRd, decreased FVFi and FVFd, and reduced neuronal surface area. Our data support the hypotheses that TRIM67 altered exocytic mode in part by regulating SNAP47 incorporation into SNARE complexes, and that SNAP47 pauses the fusion pore in an open state.

### Distinguishing exocytic modes with novel classifiers and protein perturbation

Distinguishing exocytic modes is complicated by the fast and diffraction-limited nature of vesicle fusion and fusion pores (Albillos et al., 1997; Bao et al., 2018). In contrast to semi-automated, supervised classification algorithms (Diaz et al., 2010; Yuan et al., 2015), our classifier requires no *a priori* information. Instead automated exocytic detection coupled with three classifiers and clustering indices repeatedly delivered the surprising finding that four classes of exocytic events exist in developing cortical neurons. This three pronged approach was robust: altering the half-life of a subset of events with HEPES did not alter classification or frequency of different exocytic modes. Further, classification was identical between VAMP2 and TfR at fusion events, even though the vesicular cargo TfR exhibited distinct diffusion dynamics in FVFi and FVFd.

Distinguishing KNR and FVF at the synapse remains controversial, relying on indirect and often population-based evidence to infer single vesicle behavior (He and Wu, 2007; Alabi and Tsien, 2013; Karatekin, 2018). Using developing neurons and their unique spatial distribution and temporal frequency of exocytosis, we resolved individual events, allowing identification of not two, but four discrete classes. Some classes exhibited a surprising delay after fusion, in which fluorescence does not decay instantaneously. This delay may occur if the fusion pore stabilized in an open state, which could facilitate the release of larger cargo. This is supported by the expression of a VAMP2 peptide known to stabilize the fusion pore (Guček et al., 2016). Alternatively, the delay may represent a “decision point” for the fusion state, prior to the vesicle undergoing FVF or KNR. Finally, the delay could occur due to an inefficiency in fusion, as the opening and dilation of the fusion pore is an energy barrier that must be overcome by SNARE machinery (Chernomordik and Kozlov, 2008).

Previous work in mature neurons supported the existence of FVF and KNR, which influenced our interpretation of the four classes of exocytosis in developing neuron(Alabi and Tsien, 2013; Albillos et al., 1997; He and Wu, 2007; Elhamdani et al., 2006). Previous work in chromaffin cells suggested additional exocytic modes differing from FVF and KNR (Chiang et al, 2014; Shin et al, 2018; Shin et al, 2020). Exocytic events previously interpreted as KNR were argued to be omega-shrink fusion, in which a vesicle fuses and adds up to 80% of its vesicle membrane before retreating from the plasma membrane, with slowed or absent diffusion of VAMP2-pHluorin (Chiang et al, 2014). Although we cannot rule out that omega-shrink fusion occurs in developing neurons, vesicles in chromaffin cells are much larger compared to the diameter of vesicles in developing cortical neurons and fluorescent decay an order of magnitude slower in chromaffin cells compared to fusion events in developing neurons. Therefore, it is unlikely that fusion events in these two systems will be comparable.

### Exocytic mode and morphogenesis

Prior to synaptogenesis, exocytosis provides membrane material to the expanding PM (Pfenninger, 2009; Urbina et al., 2018), which would suggest FVF predominates in developing neurons. In contrast in a mature neuron, secretion via KNR may be more relevant to maintain a steady-state surface area. We found that FVFi events comprised most exocytic events in developing neurons. The increased KNR and decreased FVF that resulted in reduced surface area in *Trim67*^*-/-*^ neurons supported a role for FVF events as a mechanism for providing new membrane required for plasma membrane expansion during morphogenesis. Interestingly, TRIM67 protein levels drop in the mature cortex once morphogenesis is complete (Boyer et al., 2018). Whether KNR-like modes of exocytosis increase at these later developmental stages remains to be determined.

Simulations based on frequency and mode exocytosis, frequency of endocytosis, and respective vesicle size remarkably recapitulated developmental increases in neuronal surface area, fitting better than simulations assuming all fusion events were FVF. Altering exocytic mode or frequency in model simulations to mimic *Trim67*^*-/-*^ or *Trim9*^*-/-*^ neurons accurately predicted altered surface areas. Although the simple model makes a number of assumptions, including that vesicles fusing via different modes are the same size and that vesicles sizes and rates of endocytosis are not altered by deletion of *Trim9* or *Trim67*, the accurate fit to empirical data suggests that our model captures relevant parameters to accurately predict surface area expansion. However, simulations in *Trim9*^*-/-*^ neurons overestimated expansion, suggesting possibly that endocytosis was also elevated in these neurons. We conclude that VAMP2-mediated fusion is the primary source of membrane addition in developing neurons, and that manipulating exocytic frequency or mode alters neuronal growth. The accuracy of our model decreased somewhat at 3 DIV, suggesting additional parameters in membrane remodeling were not captured. Perhaps analogously, *in vivo Vamp2*^*-/-*^ neurons have a largely normal morphology, suggesting compensatory mechanisms such as membrane added via ER-PM contact sites or exocytosis mediated by other SNARE proteins like VAMP7 (Fuschini et al., 2018; Galli et al., 1998) participate in membrane remodeling.

### Regulation of Exocytic Mode

Regulation of fusion pore kinetics and mode of exocytosis by regulatory proteins and number of SNAREs are well-documented (Bao et al., 2018; Bretou et al., 2014; Gauthier et al., 2011; Wen et al., 2016; Logan et al., 2017; Segovia et al., 2010; Archer et al., 2002), but SNARE complex composition adds another layer of regulation. Our findings suggested that regulation of exocytic mode in developing neurons was facilitated by increased incorporation of SNAP47, a member of the SNAP25 family of Qb/Qc t-SNAREs, into SNARE complexes. SNAP47 is capable of substituting for SNAP25 to form stable SNARE complexes with VAMP2 and syntaxin-1 using purified recombinant proteins. These complexes however, are inefficient in liposomal fusion assays, suggesting that fusion is slow to proceed in comparison to SNAP25 containing SNARE complexes (Holt et al., 2006). This is consistent with our finding that increased incorporation of SNAP47 into exocytic SNARE complexes increased the delay time between fusion pore opening and fluorescence decay caused by vesicle reacidification or VAMP2-pHluorin diffusion. We hypothesize that the SNAP47-containing SNARE complexes may be the primary SNARE complex involved in KNRd. Interestingly SNAP47 has many unique features that may be responsible for altered fusion kinetics: it lacks cysteine palmitoylation-mediated membrane targeting, has a long N-terminal extension, and has a linker region between the Qb and Qc SNARE domains (Holt et al., 2006). Structure-function experiments and observations that TRIM67 interacted with and colocalized with SNAP47 distal to exocytic sites, and that SNAP47 co-localized with a subset of VAMP2-pHluorin events with distinct fusion behavior, fit with a hypothesis that TRIM67 regulated exocytic mode via a transient interaction with SNAP47. Future studies will examine the mechanisms by which TRIM67 regulates SNAP47 protein level and incorporation into SNARE complexes, and how these may be related.

SNAP47 knockdown did not completely rescue the distribution in modes of exocytosis in *Trim67*^*- /-*^ neurons, suggesting that TRIM67 regulates exocytic fusion by multiple mechanisms. Myosin II, actin remodeling at the PM, and membrane tension are all suggested to alter exocytosis (Aoki et al., 2010; Wen et al., 2016). TRIM67 interacts with the actin-polymerase VASP at the growing tips of filopodia in the growth cone, a highly dynamic region with local changes in membrane tension and actin remodeling (Boyer et al., 2020; Staykova et al., 2011; Wen et al., 2016), but whether VASP alters exocytosis is not known. There are a host of proteins that may regulate exocytic mode, such as Cdc42, α-synuclein, and synaptotagmins (Bretou et al., 2014; Gauthier et al., 2011; Wen et al., 2016; Logan et al., 2017; Segovia et al., 2010). Separate pools of vesicles with distinct fusion characteristics and cargo (Hua et al., 2011; Rizzoli and Jahn, 2007) may fuse with different modes of fusion. Future work must define how TRIM67 interfaces with these vesicle populations and distinct mechanisms regulating fusion, and whether this unique regulation occurs at the synapse or is specific to earlier developmental time points.

## Methods

### Reagents

#### Plasmids and antibodies

Plasmid encoding human-VAMP2-pHluorin was acquired from James Rothman (Yale, New Haven, CT). VAMP2-tagRFP was generated by cloning from VAMP2-pHluorin into the tagRFP plasmid (Clontech). VAMP2^1-96^-tagRFP was created by PCR of VAMP2 (AA 1-96) into a tagRFP vector. VAMP7ΔLD-GFP and Myc-SNAP47 was obtained from Thierry Galli (University of Paris, France). VAMP7ΔLDwas clonedinto pCax-pHluorin. SNAP47-tagRFP was generated by PCR of SNAP47 from Myc-SNAP47 followed by cloning into a tagRFP vector. TRIM67 constructs were generated as described in (Boyer et al., 2020). Flag-tagged Ubiquitin was obtained from Ben Philpot (University of North Carolina at Chapel Hill, Chapel Hill, NC). GFP-CAAX was obtained from Richard Cheney (University of North Carolina at Chapel Hill, Chapel Hill, NC). TfR-pHuji was obtained from David Perrais (Université de Bordeaux, France, Addgene Plasmid #61505). Antibodies included rabbit polyclonal against TRIM67 (Boyer et al., 2018); mouse monoclonal against TRIM9 (H00114088-M01, Abnova); mouse monoclonal against c-Myc (9E10); mouse monoclonal against human β-III-tubulin (TujI SCBT); ubiquitin (sc-8017, SCBT); GAPDH (sc-166545, SCBT); GFP-Trap® Agarose (Chromotek); VAMP2 (D6O1A) Rabbit mAb #13508 (Cell Signaling); Syntaxin-1 (sc-12736, Santa Cruz); SNAP25, Mouse monoclonal (Synaptic Systems 111 011); SNAP47, polyclonal rabbit (Synaptic Systems 111403); HA, from the lab of Patrick Brennwald (UNC: Chapel Hill); IRDye® 800CW Donkey anti-Mouse IgG Secondary Antibody and IRDye® 680LT Goat anti-mouse Secondary Antibody (Licor); rabbit polyclonal against HA (71-5500, Thermo Fisher Scientific); sepharose beads (Sigma) siRNAs pool targeting SNAP47 were purchased from Dharmacon (target sequence 5’-CGTACGCGGAATACTTCGA-3’).

#### Animals

All mouse lines were on a C57BL/6J background and bred at UNC with approval from the Institutional Animal Care and Use Committee. Mouse colonies were maintained in specific pathogen-free environment with 12-12 hr light and dark cycles. Timed pregnant females were obtained by placing male and female mice together in a cage overnight; the following day was designated as embryonic day 0.5 (E0.5) if the female had a vaginal plug. *Trim9*^*-/-*^ mice were described in Winkle et al., 2014 and *Trim67*^*-/-*^ mice were described in (Boyer et al., 2018). Mice were genotyped using standard procedures. For all culture experiments, embryos from time-matched homozygous WT, homozygous *Trim9*^*-/-*^, or homozygous *Trim67*^*-/-*^ were used.

#### Culture, transfection, and treatment of primary neurons and HEK293 cell lines

Male and female embryos were used to generate cultures and were randomly allocated to experimental groups. Cortical neuron cultures were prepared from day E15.5 embryos as previously described (Viesselmann et al., 2011). Briefly, cortices were micro-dissected and neurons were dissociated with 0.25% trypsin for 15 min at 37°C followed by quenching with neurobasal media supplemented with 10% FBS and 0.5 mM glutamine. Cortices were gently triturated 15x and cells were counted by hemocytometer. Cells were spun at 0.1xg for 7 min at room temperature. Pelleted neurons were resuspended in neurobasal media supplemented with B27 (1:50 of manufacturer stock (Invitrogen)) and plated on cover glass, Mattek dishes, or tissue culture plastic coated with 1 mg/ml poly-D-lysine (Sigma-Aldrich). This same media was used for all time-lapse experiments. For transfection of plasmids and siRNA, neurons were resuspended after disassociation in Lonza Nucleofector solution (VPG-1001) and electroporated with Amaxa Nucleofector according to manufacturer protocol. A pool of 4 siRNAs targeting mouse SNAP-47 and control Luciferase siRNA (Target Sequence: 5’-CGTACGCGGAATACTTCGA-3’) (Dharmacon, Thermofisher Scientific) were electroporated along with GFP into E15.5 cortical neurons (2 µg GFP + 50 pmol siRNAs). Neurons were fixed and immunostained at 3 DIV. For TeNT treatment of VAMP2-pHluoring expressing neurons, 50 nM of TeNT (Sigma) was added to the neuronal media 30 min prior to imaging. For HEPES treatment, VAMP2-pHluorin exocytic events were imaged immediately after addition of 60 mM HEPES to the imaging media. For the TeNT and HEPES treatment, neurons were electroporated with VAMP2-pHluorin and imaged at 2 DIV. as described in “Live imaging and image analysis”. For assays investigating SNAP47 protein stability, 2 DIV neurons were treated with cycloheximide (50 μg/μl) for 0, 2, 4, 8 hrs or cycloheximide (50 μg/μl) and bortezomib (200 nM) for 8 hrs or cycloheximide (50 μg/μl) and chloroquine (50 μM) for 8 hrs prior to lysis.

HEK293 cells (female) were obtained from Rothenfußer (Klinikum der Universität München, München, Germany). HEK293 cells were maintained at 5% CO_2_ at 37°C in DMEM with glutamine (Invitrogen) supplemented with 10% FBS (Hyclone). *TRIM67*^*-/-*^ HEK293 cells were generated from this line by CRISPR/Cas9 editing as described in (Boyer et al., 2020). HEK293 cells were transfected with Lipofectamine 2000 (Invitrogen) or Polyplus jetPRIME® reagent according to manufacturer protocol.

### Imaging

#### Live and fixed cell imaging

All live cell time-lapse imaging was performed on an inverted Olympus microscope (IX81-ZDC2) with MetaMorph acquisition software, an Andor electron-multiplying charge-coupled device (iXon), and a live cell imaging chamber (Tokai Hit Stage Top Incubator INUG2-FSX). A UAPON 100x/1.49 NA DIC TIRF objective (Olympus) was used for all live cell TIRF imaging assays. GFP-CAAX images of basal surface area were acquired with an Olympus 40x, 1.4 NA Plan Apo DIC objective. The live cell imaging chamber-maintained humidity, temperature at 37°C, and CO_2_ at 5%. Fixed cells were imaged using Zeiss LSM 780 confocal laser scanning microscope using a 20x/0.75 U-Plan S-Apo objective at room temperature in glycerol and n-propyl-gallate-based mounting medium.

For all exocytosis assays, E15.5 murine cortical neurons expressing VAMP2-pHluorin, VAMP7ΔLD-pHluorin, or Tfr-pHuji and/or tagRFP expressing constructs were imaged at the 2 DIV with a 100x 1.49 NA TIRF objective and a solid-state 491-nm laser illumination and a solid-state 561-nm laser, both at 35% power at 110-nm penetration depth, unless indicated otherwise. Images were acquired using stream acquisition, imaging at 10 Hz for 2 min with 100 ms exposure time. For all colocalization experiments and dual-color live cell imaging, a Hamamatsu W-VIEW GEMINI image splitter optic (A12801-01) was used for simultaneous imaging of both lasers by splitting and projecting the beams, by wavelength, side-by-side onto an electron-multiplying charge-coupled device (iXon).

### Image analysis

ImageJ software was used for viewing of images and frames.

#### Detection of exocytic events

To identify the soma and neurites, the soma was segmented from the neurites as a rough ellipsoid encompassing the body of the neuron. All VAMP2-pHluorin, tfR-pHuji, and VAMP7ΔLD-pHluorin exocytic events were detected using the automated detection software as described (Urbina et al., 2018) in Matlab and R with Rstudio. Briefly, the algorithm defines exocytic events as transient, non-motile (mean-square displacement <0.2 μm^2^), gaussian shaped objects that reached peak fluorescent intensity four deviations above the local background intensity.

#### Analysis of exocytic events

Once events were detected, a region of interest (ROI) of 25×25 pixels surrounding the gaussian-shaped exocytic event and the bordering plasma membrane 10 frames before (1 second) and 100 frames following (10 seconds) was used for subsequent event analysis. The border plasma membrane was defined as follows: A Difference-of-Gaussian (Marr and Hildreth, 1979) filter was performed on each ROI around an exocytic event to segment the gaussian shaped fluorescent signal at timepoint 1. The border pixels of exocytic events were defined as the average fluorescence of a 15 pixel-wide perimeter surrounding the segmented gaussian shaped fluorescence.

Several parameters were measured within this ROI (Table S2). This includes the normalized change in fluorescence (ΔF/F) of the gaussian, defined as the (average intensity of an ROI – background fluorescence)/background fluorescence. The background fluorescence was calculated as the average of the region of the gaussian in the first 10 frames (1 second) before an exocytic event is detected. To ensure all relevant fusion information was obtained, fluorescence was tracked for 10 seconds following an exocytic event because a subset of exocytic events have an extended decay; this empirically determined time-frame ensured fluorescence returned to baseline (Urbina et al., 2018).

To determine the rate of decay of fluorescence, an exponential decay model was used for FVFi and KNRi:

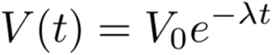

The half-life (t_1/2_ (λ)) was estimated by fitting a linear model to the log-fluorescence of the guassian-shaped exocytic event from from peak ΔF/F until the end of decay.

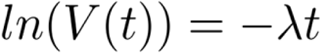

In which V(t) is the fluorescence intensity at each timepoint and t is time in seconds.

To estimate the rate of decay of FVFd and KNRd, two sequential exponential decay models were fit to the gaussian. The log of fluorescence from peak ΔF/F until the end of decay were fit using segmented regression, with the breakpoint between the two regressions decided by iteratively moving the breakpoint one time-step, fitting segmented regression, and minimizing the sum of squared error. The breakpoint that has the lowest total error was considered the true breakpoint. The first regression represents the “delay” before decay starts, and second regression’s t_1/2_ represents the fluorescence decay.

### Classificationof exocytosis

Exocytic events were classified using three independent methods of classification: Feature extraction and principle component analysis, hierarchical clustering, and dynamic time warping (DTW).

#### Feature selection and principal component analysis

Feature selection was performed using both Matlab and R. In Matlab, image features were extracted from the ROI. In R, features were extracted from the the gaussian shaped exocytic event, and the bordering plasma membrane (example in **Fig 1B**). A full list of features extracted is appended in Table S2. After features were extracted from all exocytic events, principal component analysis was performed using the R package “FactoMineR” (Lê et al. 2008) using the singular value decomposition approach. The principal components that captured the top 85% of variance were kept and used for the committee of indices.

#### Hierarchical clustering

Hierarchical clustering was performed using R’s base “stats” package and the “cluster” package.

##### Agglomerative

This function performed a hierarchical cluster analysis using a set of dissimilarities for the n objects being clustered. Initially, each object was assigned to its own cluster and then the algorithm proceeds iteratively, at each stage joining the two most similar clusters, continuing until there was a single cluster. At each stage distances between clusters were recomputed by the Lance–Williams dissimilarity update formula. In hierarchical cluster displays, a decision was needed at each merge to specify which subtree should go on the left and which on the right. Since, for n observations there are n−1 merges, there are 2(n−1) possible orderings for the leaves in a cluster tree, or dendrogram. The algorithm iordered the subtree so that the tighter cluster was on the left (the last, i.e., most recent, merge of the left subtree was at a lower value than the last merge of the right subtree). Single observations were the tightest clusters possible, and merges involving two observations were ordered by observation sequence number.

##### Divisive

fully described in chapter 6 of Kaufman and Rousseeuw (1990). The algorithm constructed a hierarchy of clusterings, starting with one large cluster containing all n observations. Clusters were divided until each cluster contains only a single observation. At each stage, the cluster with the largest diameter were selected. (The diameter of a cluster was the largest dissimilarity between any two observations.) To divide the selected cluster, the algorithm first selected the most disparate observation (i.e., the largest average dissimilarity to the other observations of the selected cluster). This observation initiated the “splinter group”. In subsequent steps, the algorithm reassigned observations that were closer to the “splinter group” than to the “old party”. The result was a division of the selected cluster into two new clusters.

#### Dynamic Time Warp

Dynamic time warping was implemented using the ‘dtw’ package in R (Giorgino, 2009) to compare two time series, X = (x_1_,…, x_N_) and Y = (y_1_,…, y_M_). DTW assumed a nonnegative, local dissimilarity function f between any pair of elements x_i_ and y_j_, with the shortcut:

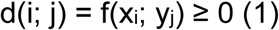

d, the cross-distance matrix between vectors X and Y, was the only input to the DTW algorithm. The warping functions φx and φy remapped the time indices of X and Y respectively. Given φ, the average accumulated distortion between the warped time series X and Y was computed:

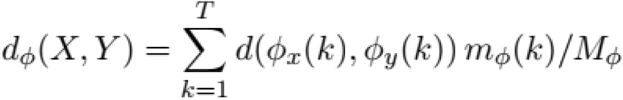

where mφ(k) was a per-step weighting coefficient and Mφ was the corresponding normalization constant, which ensured that the accumulated distortions were comparable along different paths. To ensure reasonable warps, constraints were imposed on φ. Monotonicity was imposed to preserve their time ordering and avoid meaningless loops:

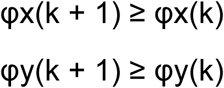

The goal of DTW was to find the optimal alignment φ such that

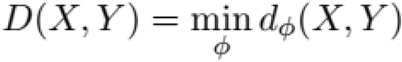

Which represented the warp path cost of X and Y. The warp path cost for each exocytic event pairing was built into a matrix.

#### Clustering

Each of the classifiers were independently clustered by a committee of indices. Numerous clustering validity algorithms have been proposed (Rousseeuw, 1987; Pelleg et al., 2000; Sugar and James, 2003). These algorithms combine information about intracluster compactness and intercluster isolation, as well as other factors, such as geometric or statistical properties of the data, the number of data objects, and the dissimilarity or similarity measurements. However, different algorithms lead to different clusters, and even in a single algorithm, choice of parameters can lead to different clusters. To perform classification, a plurality-rules decision of 20 well-used algorithms for determining clustering was used. This committee of indices includedA full list of the 20 methods used in this committee of indices can be found in Charrad et al, 2014.

The analysis described above was developed for VAMP2-pHluorin, and exploited for VAMP7ΔLD-pHluorin and TfR-pHuji. Adjustments were made in TfR-pHuji analysis because it was larger (∼980 amino acids) than VAMP2-pHluorin (∼350 amino acids) and had distinct diffusion characteristics (Di Rienzo et al., 2013; Fujiwara et al., 2002; Fujiwara et al., 2016) TfR undergoes hop diffusion, a model of transmembrane protein diffusion characterized by various transmembrane proteins anchored to the actin-based membrane skeleton meshwork acting as rows of pickets that temporarily confine diffusion. To capture this information, an additional parameter measuring goodness-of-fit to linear or exponential decay was added. The goodness-of-fit was defined as the R^2^ of fitting a linear regression to the fluorescent decay divided by the R^2^ of fitting an exponential curve for the half-life as described under “analysis of exocytic events”. A linear fit will have a high linear goodness-of-fit, while an exponential curve will have a lower value.

#### Colocalization

To perform co-localization analysis, TRIM67tagRFP and SNAP47tagRFP puncta were identified by difference-of-gaussian filter. The x,y centroids of VAMP2-pHluorin events, SNAP47-tagRFP puncta, or TRIM67-tagRFP puncta were fit to a marked point process model, which allowed comparisons of groups of points, and Ripley’s L value distances were measured.(Grantham, 2012). Ripley’s L value measures the average distance between points normalized to the number of puncta and total cellular area. This average distance was statistically compared to what distances expected if points were “randomly” localized, allowing a measure of colocalization. Ripley’s L values here were compared to theoretically random marked points, setting a threshold of 0.05% chance to fall below the theoretical Ripley’s L value. All values were normalized to the theoretical Ripley’s L value for comparisons.

### Surface area measurements and modeling predictions

#### Empirical Surface Area Estimation

To estimate the surface area of neurons during DIV 1-3, TIRF images of GFP-CAAX expressing cells at indicated time points were used. The soma was identified and segmented from the neurites, and the basal surface area of each were calculated separately. Total neurite area was estimated by doubling the measured basal membrane area. The average height of the soma (∼11 μm) was obtained from confocal image z stacks through cortical neurons at 2 DIV. The major and minor axis of the soma were measured, and used with the average height of the soma to calculate the surface area using the following calculation for a truncated ellipsoid:

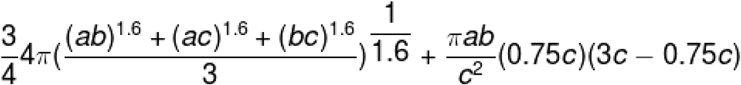

Where a,b, and c represent the 3 axes of the soma (length, width, and height, respectively). The surface area of soma and neurites were summated to obtain the final surface area estimate of the neuron.

#### Model Selection

A Bayesian linear model based on Urbina et al., 2018 was adapted to predict neuronal surface area increases from DIV1 to DIV 2 and 3. Surface area at 2 and 3 DIV, the predicted and independent variables, are a positive, continuous, right skewed distribution, which has non-uniform standard deviation over time. Several models were constructed, selecting for probability distributions that would fit the criteria of positive, continuous, and right-skewed data: log-normal, Skew-normal, and Gamma. Models using the t-distribution and the normal distribution as a base model were also constructed, based on their well-characterized distributions. Previously published data were used as priors. Namely, the surface area of neurons at 1 DIV, the surface area of vesicles, the frequency of exocytosis, the frequency of clathrin-mediated endocytosis, and the size of clathrin-coated vesicles were used. This was combined with newly measured surface area of neurons at 1 DIV (described under “Surface area estimation”) to model membrane addition. This provided confidently constructed priors that were estimated from previously measured neuronal surface area, instead of subjectively chosen priors. From this, four hierarchical models of surface area expansion were constructed: Gamma with a log-link, log-normal, skewed-normal, and log-normal with a lower bound truncation of 0. Given the difference in the standard deviation between the observed surface area at 1 and 2 DIV (Urbina et al., 2018), a fifth model was added, in which the standard deviation was also modeled based on the measured standard deviation at 1 and 2 DIV. Four chains were run with 100,000 samples after a burn-in period of 10,000 steps for a total of 110,000 steps to ensure chain convergence and enough coverage of the posterior distribution. Model evaluation using Rhat and ESS suggested the chains were well-mixed and converged properly. Each of these models were then compared by computing Watanabe-Akaike information criteria and using leave-one-out validation to compute the expected log pointwise predictive distribution for the difference in each of the models. Model selection using these criteria indicate that using the log-normal family results in the best fit for these data, followed by the gamma distribution. Taking all of these terms into account, an equation for the hierarchical Bayesian linear model was created, using the log-normal distribution:

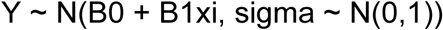

### Western blot

#### Analysis of protein lysates

To investigate SNAP47, SNAP25, and GAPDH protein levels, cultured E15.5 cortical neurons of indicated genotypes were lysed in immunoprecipitation (IP) buffer (10% glycerol, 1% NP-40, 50 mM Tris pH 7.5, 200 mM NaCl, 2 mM MgCl_2,_ 1 mM NEM, and protease and phosphatase inhibitors at 2 DIV. Lysates were collected and centrifuged at 19,745 xg for 10 min. Supernatants were diluted in 4x sample buffer (200 mM Tris-Cl, pH 6.8, 8% SDS, 0.4% Bromophenol Blue, 40% glycerol, 20% 2-Mercaptoethanol), boiled for 10 min at 100 °C, and resolved by SDS-PAGE. For SNAP47 half-life measurements, cycloheximide treated cortical neurons were lysed (300 mM sucrose, 1% NP-40, 50 mM Tris pH 7.5, 200 mM NaCl, 2 mM MgCl_2_ and protease, phosphatase and deubiquitinase inhibitors). Lysates were collected and centrifuged at 19,745 xg for 10 minutes. Protein concentrations were estimated, and lysates were boiled with 2x sample buffer and resolved by SDS-PAGE.

#### Co-immunoprecipitation assay

For coimmunoprecipitation assays in HEK293 cells expressing GFP-SNAP47 and either MycTRIM9, MycTRIM67, or MycTRIM67 domain deletion mutants. were lysed in IP buffer and lysates collected and centrifuged at 19745xg for 10 min. 500–1,000 µg of protein per sample was used per IP (the same amount of protein was maintained across all conditions per experiment). Myc-tagged proteins were precipitated with anti-Myc antibody (9E10) overnight, followed by Protein A coupled agarose beads. Beads were washed twice with IP buffer, pelleted, and boiled with 2X sample buffer. For endogenous SNAP47 immunoprecipitation from cultured cortical neurons, E15.5 cortical neurons at 2 DIV were treated with 1mM NEM (15 minutes) prior to lysis. Membrane proteins were enriched through ultra-centrifugation of neuronal lysates, as described in (Shimojo et al., 2015). Briefly, cortices of wild-type mice were homogenized in extraction buffer (20 mM HEPES-NaOH (pH 7.4), 320 mM sucrose, 5 μg/ml leupeptin, 2 μg/ml aprotinin, and 1 μg/ml pepstatin) post NEM treatment. Membrane fractions were isolated by two sequential steps of centrifugation at 3,000 and 100,000 × g and solubilized for 30 min in 20 mM HEPES-NaOH (pH 7.4), 150 mM NaCl, and 1% Triton X-100. Protein extracts were cleared by centrifugation at 100,000 × g and incubated overnight with sepharose beads coated with anti-SNAP47, or control IgG antibodies. Attached complexes were then washed five times with extraction buffer, eluted from the beads with an SDS gel loading buffer, and resolved by SDS-PAGE.

#### SNARE complex assay

SNARE complex assays were performed as described in (Hayashi et al., 1994) with modification. E15.5 cortical neurons at 2 DIV were treated with either 1mM NEM (15 minutes) or 1mM NEM+ 2mM DTT (15 minutes) prior to lysis with homogenization buffer (10mM HEPES-NaOH, pH 7.4, 150mM NaCl, 1mM EGTA, 1mM NEM, and protease inhibitors). After collection of lysates, Triton X-100 was added to a final composition of 1% Triton x-100 followed by trituration 10x and solubilization for 2 min on ice. Lysates were then centrifuged at 10min, 6000 g. Samples were diluted in 4x sample buffer (200mM Tris-Cl, pH 6.8, 8% SDS, 0.4% Bromophenol Blue, 40% glycerol, 20% 2-Mercaptoethanol) into two replicates, one replicate incubated at 100 °C for 10 min and the second incubated at 37 °C for 10 min. SNARE complexes and monomers were then resolved by SDS-PAGE.

#### Ubiquitination experiment

For ubiquitination assay, *TRIM67*^*-/-*^ or rescued (transfected with Myc or Myc-TRIM67 respectively) HEK cells were co-transfected with HA-tagged Ubiquitin and control GFP or GFP-SNAP47 using Lipofectamine 2000. GFP-Trap beads (Chromotek) were used to enrich GFP-SNAP47 and GFP according to manufacturer protocol. Briefly, cells were treated with MG132 for 4 h then lysed in ubiquitination immunoprecipitation (Ub-IP) buffer (50 mM Tris-Cl, 150 mM NaCl, 1 mM EDTA, 0.5% Triton X, 0.7% *N*-ethylmaleimide, and protease and phosphatase inhibitors, pH 7.3–7.4). Lysates were collected and centrifuged at 19745 xg for 10 min 4°C. 500–1,000 µg of protein per sample was used in each assay (the same amount of protein was maintained across all conditions per experiment). 15 µl GFP-TRAP beads were incubated with the lysate for 2.5 hrs followed by three washes. The first wash was performed using 10mM Tris-Cl, pH7.5, 150 mM NaCl, 0.5 mM EDTA, 0.7% NEM. This was followed by two washes using first a stringent buffer (8 M Urea, 1% SDS) and then SDS wash buffer (1% SDS). The beads were then boiled with 2X sample buffer. HA-ubiquitin that co-immunoprecipitated with GFP-SNAP47 and migrated at a higher apparent molecular weight was interpreted as ubiquitinated SNAP47 (SNAP47-Ub)

#### Immunoblotting

All protein samples were resolved by SDS-PAGE followed by transfer onto 0.45μM (0.22μM for the SNARE complex assay) nitrocellulose paper and analyzed by immunoblotting. Blots were probed with indicated primary antibodies, followed by IRDye® 680LT Goat anti-Rabbit Secondary Antibody and/or IRDye® 800CW Donkey anti-Mouse IgG Secondary Antibody and imaged on an Odyssey Licor. Western blots were analyzed using Fiji (ImageJ) or Li-Cor Image Studio Lite. Total fluorescence of labeled bands representing relative protein were normalized to control conditions for each experiment as indicated. For all western blot comparisons, the relative band intensity or relative ratios (indicated for each blot) were log-transformed followed by paired-t-test..

### Mass Spectrometry

To identify potential interacting partners for TRIM67, we performed three biological replicates using the Proximity-Dependent Biotin Identification (BioID) approach (Roux et al. JCB 2012). Briefly, the negative control Myc-BirA* and Myc-BirA* TRIM67ΔRING were packaged into Short Term Herpes Simplex Virus (HSV)(MGH) and driven under a IE 4/5 promoter. GFP expression is driven in tandem downstream of a mCMV promoter. Cortical neurons from wildtype and *Trim67*^*-/-*^ E15.5 litters were dissociated and plated on PDL coated tissue culture dishes. Approximately 40 hours post-plating the neurons were transfected with HSV (MOI = 1.0) carrying either the negative control Myc-BirA* or Myc-BirA* TRIM67ΔRING. 6 hrs post-infection neurons were treated with 50 µM Biotin for 24 hrs. After incubation the cells were lysed using RIPA buffer (150 mM NaCl, 25 mM Tris-HCl, pH 7.5, 0.1% SDS, 1.0% NP-40, 0.25% Deoxycholic acid, 2 mM EDTA, 10% glycerol, protease and phosphatase inhibitors). Biotinylated proteins were then enriched from the lysate using streptavidin-conjugated sepharose beads. The enriched proteins were digested with trypsin and eluted using the RapiGEST SF Surfactant protocol (Waters). C18 column and Ethyl acetate extraction protocols were employed to prepare the peptides for mass spectrometry.

Reverse-phase nano-high-performance liquid chromatography (nano-HPLC) coupled with a nanoACQUITY ultraperformance liquid chromatography (UPLC) system (Waters Corporation; Milford, MA) was used to separate trypsinized peptides. Trapping and separation of peptides were performed in a 2 cm column (Pepmap 100; 3-m particle size and 100-Å pore size), and a 25-cm EASYspray analytical column (75-m inside diameter [i.d.], 2.0-m C18 particle size, and 100-Å pore size) at 300 nL/min and 35°C, respectively. Analysis of a 150 min gradient of 2% to 25% buffer B (0.1% formic acid in acetonitrile) was performed on an Orbitrap Fusion Lumos mass spectrometer (Thermo Scientific). The ion source was operated at 2.4kV and the ion transfer tube was set to 300°C. Full MS scans (350-2000 m/z) were analyzed in the Orbitrap at a resolution of 120,000 and 1e6 AGC target. The MS2 spectra were collected using a 1.6 m/z isolation width and were analyzed either by the Orbitrap or the linear ion trap depending on peak charge and intensity using a 3 s TopSpeed CHOPIN method.32 Orbitrap MS2 scans were acquired at 7500 resolution, with a 5e4 AGC, and 22 ms maximum injection time after HCD fragmentation with a normalized energy of 30%. Rapid linear ion trap MS2 scans were acquired using an 4e3 AGC, 250 ms maximum injection time after CID 30 fragmentation. Precursor ions were chosen based on intensity thresholds (>1e3) from the full scan as well as on charge states (2-7) with a 30-s dynamic exclusion window. Polysiloxane 371.10124 was used as the lock mass. All raw mass spectrometry data were searched using MaxQuant version 1.5.7.4. Search parameters were as follows: UniProtKB/Swiss-Prot human canonical sequence database (downloaded 1 Feb 2017), static carbamidomethyl cysteine modification, specific trypsin digestion with up to two missed cleavages, variable protein N-terminal acetylation and methionine oxidation, match between runs, and label-free quantification (LFQ) with a minimum ratio count of 2. To rank candidate protein-protein interactions by likelihood of interaction, LFQ values for proteins identified in control and pulldown experiments from three biological replicates were input into SAINTq (version 0.0.4). Interactions were sorted by SAINT’s interaction probability and a false discovery rate threshold of 25% was employed. A full list of identified proteins will be reported elsewhere (Menon et al, in preparation).

### Statistics and data presentation

The software package R was used for statistical analysis of data. Both R and Adobe Illustrator were used for the generation of figures. At least three independent biological replicates were performed for each assay, often more. Sample sizes for all experiments measuring frequency were estimated using power analysis (Power = 0.8) and the estimated effect size based off of previous work in Urbina et al, 2018. For two-sample comparisons of normally distributed data, Welch’s t-test was used for two independent samples, or paired t-test for paired samples. For multiple comparisons, Welch’s or paired t-tests corrected using Benjamini-Hochberg method. For comparison of ratios for the four classes, multivariate linear regression was used, with the expected ratio of the four classes as the dependent variables and the conditions as the independent variables. For analysis of Fig 1G, a chi-squared expected ratio test was performed to determine if the pHluorin classification assigned FVF or KNR classes different than randomly. For analysis of non-normally distributed data, the was used to determine significance followed by the method described above.

#### Plots

All boxplots were graphed as follows: the box represents the median value (middle of boxplot) and the interquartile range, which is the distance between the first and third quartile of the data (with the first quartile being the middle value between smallest number and median of the dataset, and the third quartile representing the middle value between the largest number and median of the dataset). The whiskers extend out to the last data point within 1.5x the interquartile range.

## Acknowledgements

The authors would like to thank Shawn Gomez for critical suggestions and comments on dynamic time warping, Caroline Monkiewicz, Vong Thoong, and Chris Hardie for mouse breeding, husbandry, and genotyping. We thank Michelle Itano and the UNC Neuroscience Imaging Core for training and access to the Zeiss 780 Laser Scanning Confocal Microscope, funded in part by P30 NS045892 and U54 HD079124. Funding from the National Institutes of Health supported this research: including R01NS112326 (SLG), R35GM135160 (SLG), and F31NS103586 (FLU).

---

**Figure S1:**
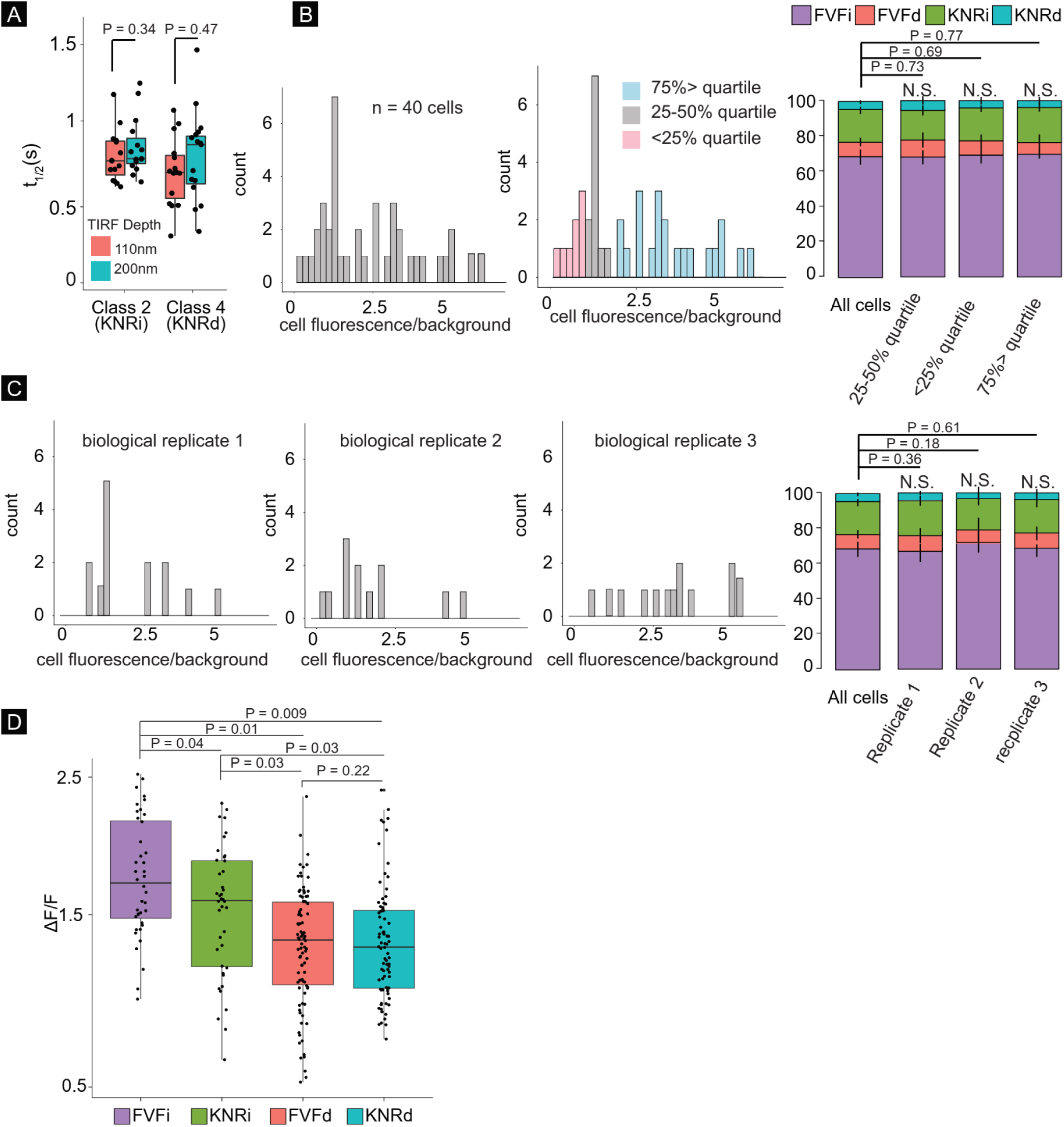
Distinguishing features of four modes of exocytosis. **A)** t_1/2_ of VAMP2-phluorin fluorescent decay in KNRi and KNRd events is the same at 110 nm and 200 nm evanescent wave TIRF penetration depth, suggesting fluorescence decay is not caused by vesicle retreat from the PM. **B)** Cellular VAMP2-pHluorin fluorescence or **C)** biological replicate did not affect exocytic mode distribution. No significant difference in mode of exocytosis was found when partitioning cells based on fluorescent signal or grouping by biological replicate. **D)** Peak ΔF/F of fusion events of each exocytic mode (t-test with Benjamini-Hochberg correction).

**Figure S2:**
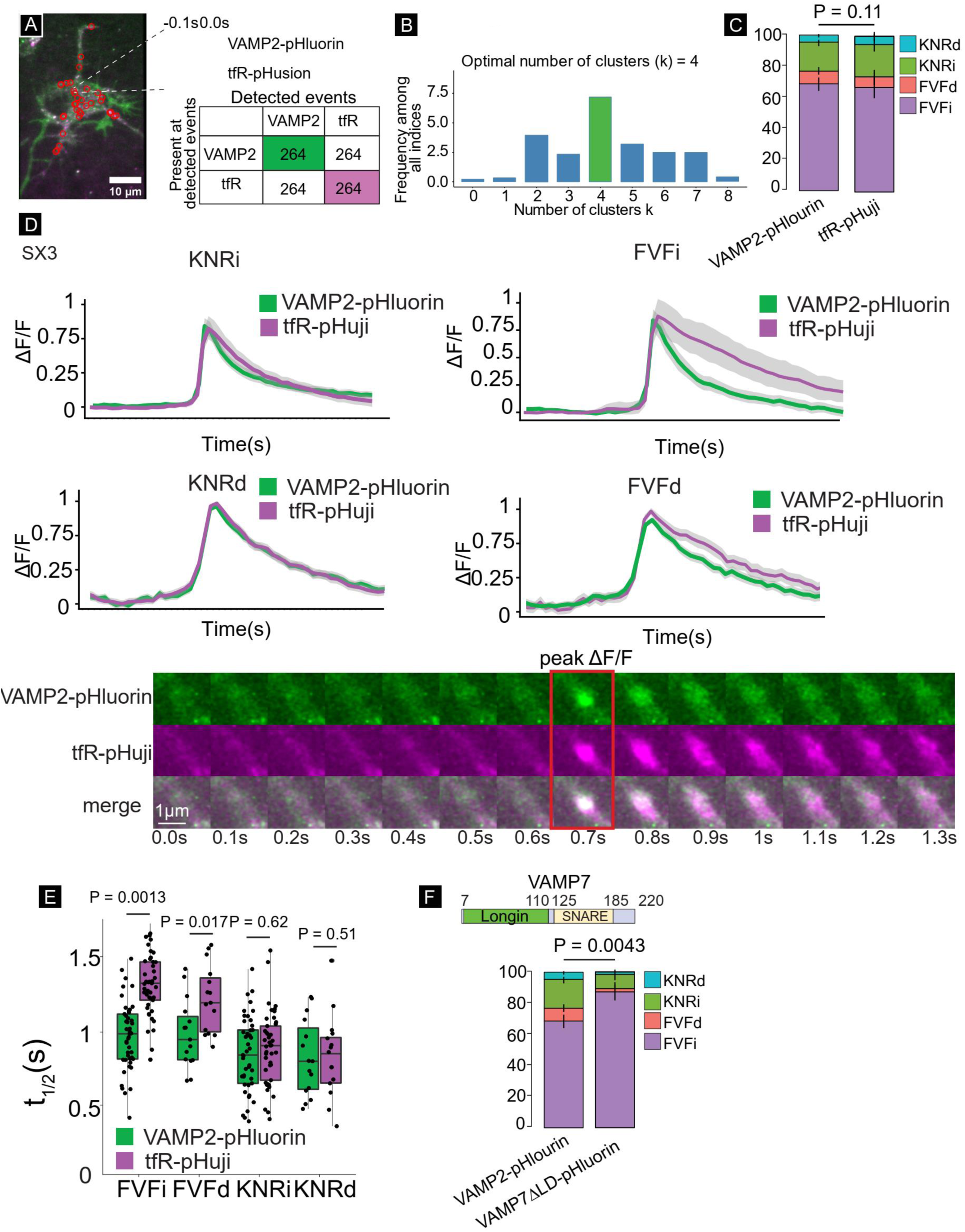
Four exocytic modes are also detected with TfR-pHuji and VAMP7ΔLD-pHluorin. **A)** Colocalization of VAMP2-pHluorin and TfR-pHuji. 100% of VAMP2-pHluorin detected events colocalized with TfR-pHuji events, and 100% of TfR-pHuji detected events colocalized with VAMP2-pHluorin (n = 6 cells, n = 264 events, white scale bar in inset = 1μm). **B)** Decision histogram of the committee of indices for TfR-pHuji, with the plurality of indices choosing four classes. **C)** Proportion of FVFi, FVFd, KNRi, and KNRd in wildtype neurons expressing VAMP2-pHluorin or tfR-pHuji (n = 11 neurons, 4 biological replicates, multivariate linear regression). **D)** Mean fluorescent curves +/- SEM (grey) of each class of VAMP2-pHluorin and TfR-pHuji. (lower) representative images of ad VAMP2-pHluorin exocytic event with TfR-pHuji. **E)** Half-life of fluorescence decay (t_1/2_) of each class of VAMP2-pHluorin (green) and TfR-pHuji (n = 11 cell, n = 4 biological replicates, paired t-tests with Benjamini-Hochberg correction) **F)** Proportion of FVFi, FVFd, KNRi, and KNRd in wildtype neurons expressing VAMP2-pHluorin or VAMP7ΔLD-pHluorin (n = 8 cells per condition, n = 3 biological replicates, multivariate linear regression)

**Figure S3:**
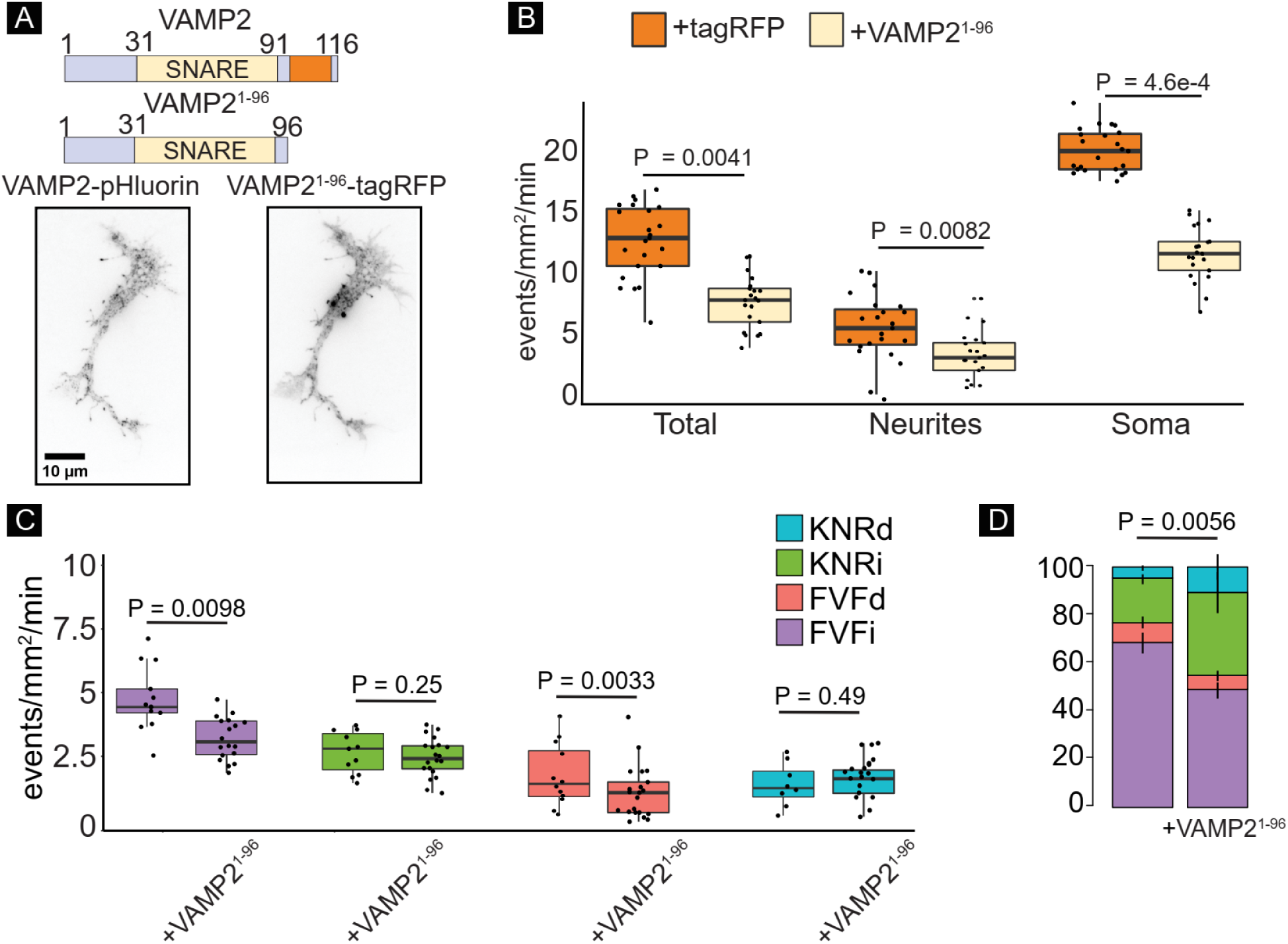
Expression of truncated VAMP2 alters exocytic mode. **A)** VAMP2 and VAMP2^1-96^ constructs and representative images of a neuron expressing both of VAMP2-pHluorin and VAMP2^1-96^-tagRFP. **B)** Frequency of exocytic events in the whole cell, soma, and neurites of neurons expressing VAMP2-pHluorin +/- VAMP2^1-96^-tagRFP (n = 23 cells per condition; n = 3 biological replicates; Welch’s t-test followed by Benjamini-Hochberg correction). **C)** Frequency of exocytic events of each class of in whole neurons expressing VAMP2-pHluorin +/- VAMP2^1-96^-tagRFP (n = 17 cells per condition; Welch’s t-test followed by Benjamini Hochberg correction). **D)** Relative proportions of each class of exocytosis in whole neurons expressing VAMP2-pHluorin +/- VAMP2^1-96^-tagRFP (n = 17 cells per condition; multivariate linear regression).

**Figure S4:**
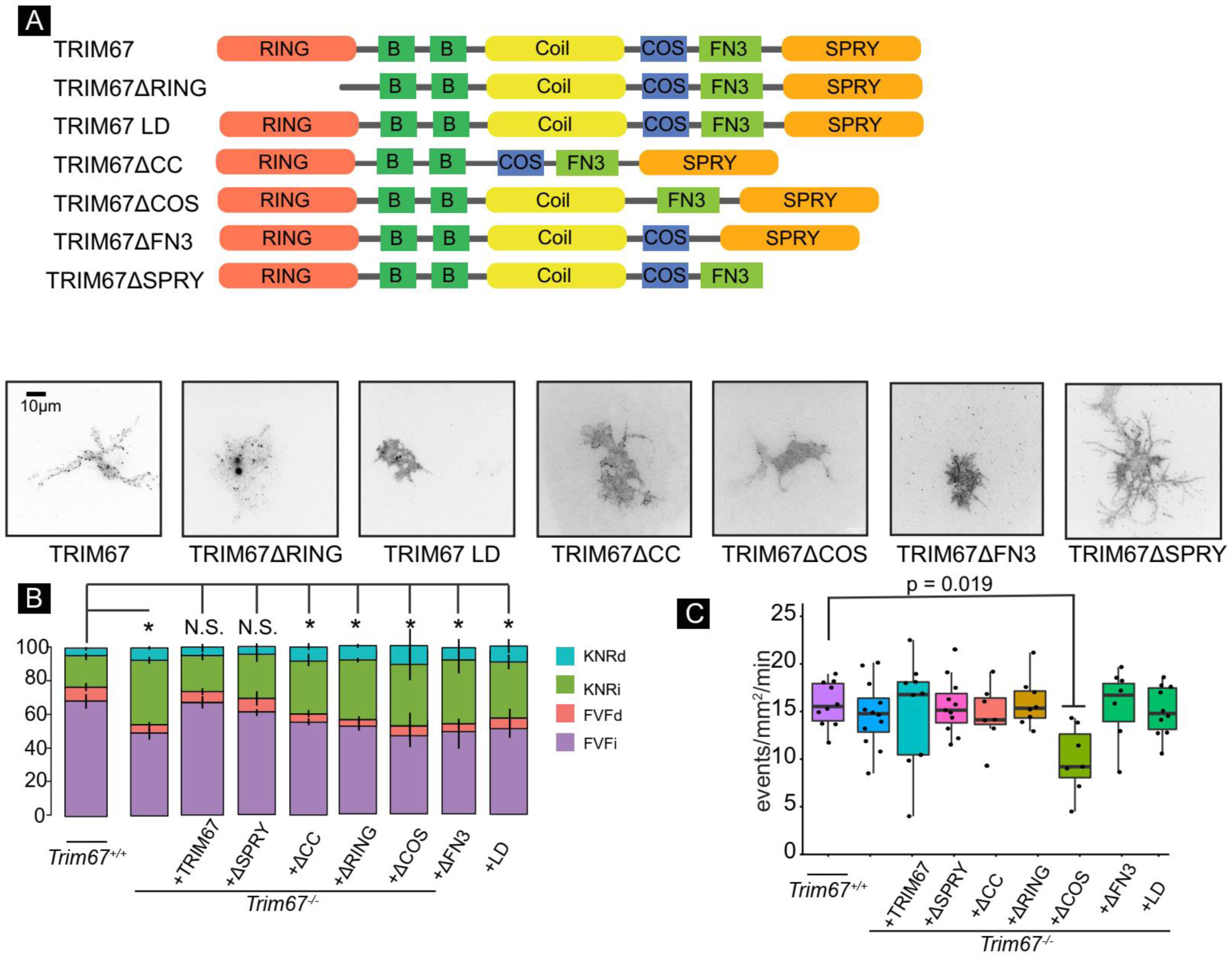
Multiple domains of TRIM67 and ligase function are required for exocytic mode regulation. **A)** Domain architecture of TRIM67 constructs tagged with TagRFP for structure:function assays. TRIM67 is a TRIpartite Motif (TRIM) E3 ubiquitin ligase, characterized by a ubiquitin ligase RING domain, two BBox domains, a coiled-coil motif (CC) that mediates multimerization, a COS domain, FN3 domain, and SPRY domain Example fluorescence images of each mutant (below). **B)** Relative proportion of each exocytic class (n = 16 cells per condition; n = 4 biological experiments; multivariate linear regression. P-values from left to right: p = 0.0004; p = 0.49; p = 0.20; p = 0.024; p = 0.009; p = 0.029; p = 0.0073; p = 0.014) and **C)** exocytic frequency in *Trim67*^*+/+*^ neurons, *Trim67*^*-/-*^ neurons, or *Trim67*^*-/-*^ neurons expressing VAMP2-pHluorin and TRIM67-tagRFP constructs (n = 14 cells per condition; n = 3 biological replicates; l Welch’s t-test with Benjamini-Hochberg correction).

**Figure S5:**
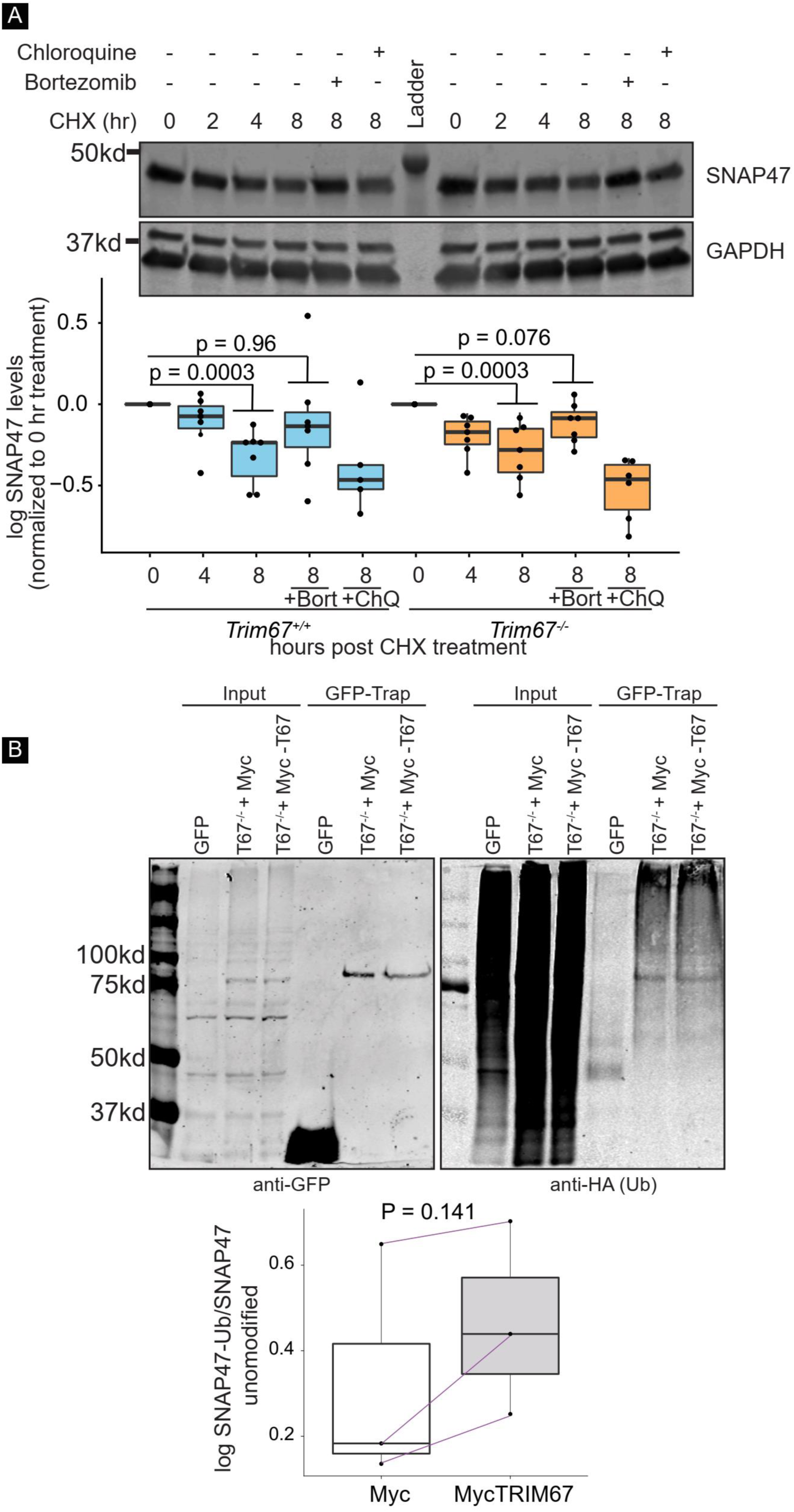
Proteasomal degradation and ubiquitination of SNAP47 are TRIM67 independent. **A)** Immunoblot of neuronal lysates at 2 DIV after treatment with cycloheximide (CHX,50 μg/μl) for 0, 2, 4, 8 hrs or cycloheximide (50 μg/μl) +/- bortezomib (Bort) (200 nM) for 8 hrs or cycloheximide (50 μg/μl) +/- chloroquine (ChQ) for 8 hours (log-ratio paired t-test (methods) performed between genotypes at each timepoint followed by Benjamini-Hochberg correction; n = 7 blots, n =5 for CHX+ChQ in *Trim67*^*+/+*^, n = 6 for CHX+ChQ in *Trim67*^*-/-*^). **B)** Immunoblot of lysates and immunoprecipitation in *TRIM67*^*-/-*^ HEK cells expressing HA-tagged ubiquitin (HA-Ub) along with empty GFP or GFP-tagged SNAP47 (GFP-SNAP47) with empty-Myc or Myc-tagged TRIM67 (Myc-TRIM67). Cells overexpressing these constructs were treated with MG132 for 4 hrs prior to lysis to prevent proteasomal degradation (log-ratio paired t-test (methods), n = 3).

## Movie Legends

**Movie 1: Heterogeneity in single vesicle fusion event kinetics**. Example time-lapse TIRF images of four classes of fusion events and surface plots (3D plots with x and y spatial axis and z intensity axis). Diffusion of VAMP2 fluorescence is evident in FVFi and FVFd, but not KNRi and KNRd, whereas the delay before onset of decay is evident in FVFd and KNRd, but not FVFi and KNRi.

**Movie 2: Expression of a truncated VAMP2 alters exocytic mode**. Time-lapse TIRF images of wildtype cortical neuron expressing VAMP2-pHluorin and VAMP2^1-96^-tagRFP at 2 DIV.

**Movie 3: TRIM67 biases exocytic mode towards full-vesicle fusion**. Time-lapse TIRF images of wildtype, *Trim9*^*-/-*^, and *Trim67*^*-/-*^ cortical neurons at 2 DIV expressing VAMP2-pHluorin.

**Movie 4: TRIM67 does not co-localize with VAMP2-pHluorin exocytic events**. Time-lapse TIRF images of a *Trim67*^*-/-*^ cortical neuron at 40 hours *in vitro* expressing VAMP2-pHluorin and GFP-TRIM67.

**Movie 5: The t-SNARE SNAP47 colocalizes with TRIM67**. Time-lapse TIRF images of a *Trim67*^*-/-*^ cortical neuron expressing GFP-TRIM67 and SNAP47-tagRFP at 2 DIV.

**Movie 6: The t-SNARE SNAP47 localizes to VAMP2-mediated exocytic events**. Time-lapse TIRF images of an automatically detected VAMP2-pHluorin containing exocytic event (green channel) with identifiable SNAP47 puncta (magenta channel), captured from a wildtype neuron expressing VAMP2-pHluorin and SNAP47-tagRFP.

## Notes

### Competing Interest Statement

The authors have declared no competing interest.

### Summary of Updates

This manuscript includes new experiments confirming the existence of four modes of exocytosis with additional markers of fusion. In addition other controls are included to confirm that VAMP2 overexpression does not induce artifacts in fusion behavior.

